# Comparative spatiotemporal single cell transcriptomes reveal rewiring of pre-existing regulations during emergence of Kranz anatomy in C_4_ grasses

**DOI:** 10.1101/2024.10.28.620769

**Authors:** Caiyao Zhao, Jianzhao Yang, Anting Ni, Hanyang Chen, Hong Su, Yating Cheng, Xiaoya Li, Liyuan Zhong, Dongming Fang, Hao-Ran Sun, Juan Yi, Ying Wang, Yanjie Wang, Ming-ju Amy Lyu, Xiaoxiang Ni, Lichuan Chen, Huanjin Li, Guichao Yu, Yumeng Wang, Rulei Chen, Hanyue Yu, Wenwen Shao, Haixi Sun, Yinqi Bai, Chuanshun Li, Yang Dong, Changsong Zou, Xiaoyu Tu, Shisong Ma, Xing Guo, Gengyun Zhang, Peng Wang, Huan Liu, Tong Wei, Ying Gu, Xun Xu, Xin-Guang Zhu

## Abstract

Many of the world’s most productive food and bioenergy crops use C_4_ photosynthesis, which have high photosynthetic efficiency due to a Kranz anatomy-based CO_2_ concentrating mechanism. Here, we took a comparative transcriptomics approach using single cell spatial transcriptomes of leaf primordia for maize (*Zea mays*) and single cell RNA-seq (scRNA-seq) atlases of corresponding leaf tissues for three C_4_ species (*Zea mays*, *Sorghum bicolor*, *Setaria viridis*) and one C_3_ rice (*Oryza sativa*) to study regulatory networks involved in the development and evolution of Kranz anatomy. We show that the formation of Kranz anatomy involves extensive recruitment and modification of pre-existing regulatory modules, especially the SHR-SCR module and the auxin signaling pathway. Members of the *INDETERMINATE DOMAIN* (*IDD*) family transcription factors, including *IDD7* and *IDDP1*, contribute to the modification of the involved modules. These extensive recruitment and modification of existing genetic regulatory programs underlie the repeated emergence of C_4_ photosynthesis during evolution.

## Introduction

Given the high light, nitrogen and water use efficiency, C_4_ plants hold great potential in protecting global food security.^1^ C_4_ plants have evolved independently over 60 times historically.^2^ Understanding the mechanisms of C_4_ plant development and evolution could potentially enable the engineering of C_4_ photosynthesis in C_3_ crops.^3^ The superior photosynthetic properties of C_4_ plants are primarily due to a CO_2_ concentrating mechanism based on Kranz anatomy, which facilitates the compartmentalized metabolism in bundle sheath cell (BSC) and mesophyll cell (MC). So far, metabolic structure and morphological features associated with C_4_ photosynthesis are well established.^3,4^ In contrast, regulatory mechanisms underlying these different properties are largely unknown, although a number of regulators controlling Kranz anatomy have been identified, such as *GOLDEN2* (*G2*), *SHOOT ROOT1* (*SHR1*), *SCARECROW1* (*SCR1*), *TOO MANY LATERALS1* (*TML1*)) through traditional forward genetics approaches. ^5–12^

High-throughput RNA-seq methods emerged as a promising approach to reveal the mechanisms controlling Kranz anatomy development.^13–29^ For example, Wang et al (2013, 2017) performed comparative analysis on different leaf primordia of maize and identified *GLK* as a major regulator of BSC development, which might have facilitated the transition from C_3_ to C_3_-C_4_ transition.^18,30^ Camo-Escobar et al. (2024) used scRNA-seq to show a significant reduction in the proportion of BSC and a downregulation of the C_4_ shuttle system in the *shr2* mutant.^24^ By using multiplexed in situ hybridization, Perico et al. (2024) showed the spatiotemporal expression dynamics of auxin- and other vein-related genes during the development of different-rank veins in the maize leaf, suggesting auxin plays a crucial role during Kranz development.^22^ Given that the development of Kranz anatomy is highly coordinated between different cells types in space, spatial-temporal single-cell transcriptomes of leaf primordia, in contrast to the earlier whole-tissue based transcriptomics approach, hold a great promise to reveal molecular mechanisms controlling morphogenesis of the Kranz anatomy in C_4_ leaves.

In this study, we generated spatial enhanced resolution omics-sequencing (Stereo-seq) and scRNA-seq atlases for leaf primordia in maize to study the development of Kranz anatomy. With these spatial scRNA atlases, we constructed a gene regulatory network for different cell types and identified regulatory cascades that control the vein initiation and Kranz anatomy development. By also collecting and analyzing scRNA-seq atlases for leaf primordia in sorghum, foxtail-grass, and rice, we identified the new regulatory mechanisms controlling features of Kranz anatomy in C_4_ grasses. Furthermore, we show that the well-known regulatory modules including the SHR-SCR module, were recruited and modified during the emergence of C_4_ photosynthesis in grasses, with the *INDETERMINATE DOMAIN* (*IDD*) family gene members involved in the altered regulation. This study shows that there is extensive rewiring the regulatory programs during the emergence of C_4_ photosynthesis in the grass family.

## Results

### Spatial transcriptome atlas for early developmental leaves of maize differentiates the subtle differences among cell types

To profile the transcriptome of cells of early stage Kranz anatomy, we conducted the spatial transcriptome of maize leaf primordia, consisting of P3-P6 on the section of base stem at the stage of 12-day-old maize seedling (Figure 1A). We extracted *in situ* transcriptome of individual cells based on their cell wall outlines stained with fluorochrome (see details in materials and methods, Figure 1B) and obtained the transcriptomes of 14,037 cells (Figures 1A and 1B). The expression patterns of known marker genes are in strong alignment with previously established knowledge (Figure 1C), confirming the reliability of the spatial transcriptome data. Within the 17 unsupervised clusters, we identified most of the cell types of maize primordium based on expression patterns of established marker genes (Figure 1D, Figure S1, including two clusters of pavement cells (classified as epidermis), two clusters of guard cells and/or subsidiary cells (GC/SC); five clusters of veins including phloem (cluster 7, C7, companion cells), xylem cells (C13), procambium (C12) located in the leaf primordium, older veins (C14) and sclerenchyma cells (C11) located in the leaf sheath (Figures S1B and 1C). There are also three clusters of dividing cells undefined, including two clusters of S-stage cells and one cluster of G2M-stage cells (Figures 1D, Figure S1C). Among these cell types, the two distinct clusters of epidermal cells are notably positioned within the inner and outer regions of the leaf primordia, respectively. This spatial segregation indicates that the nuanced differences between these two developmental stages of the epidermis can indeed be accurately differentiated.

**Figure 1:**
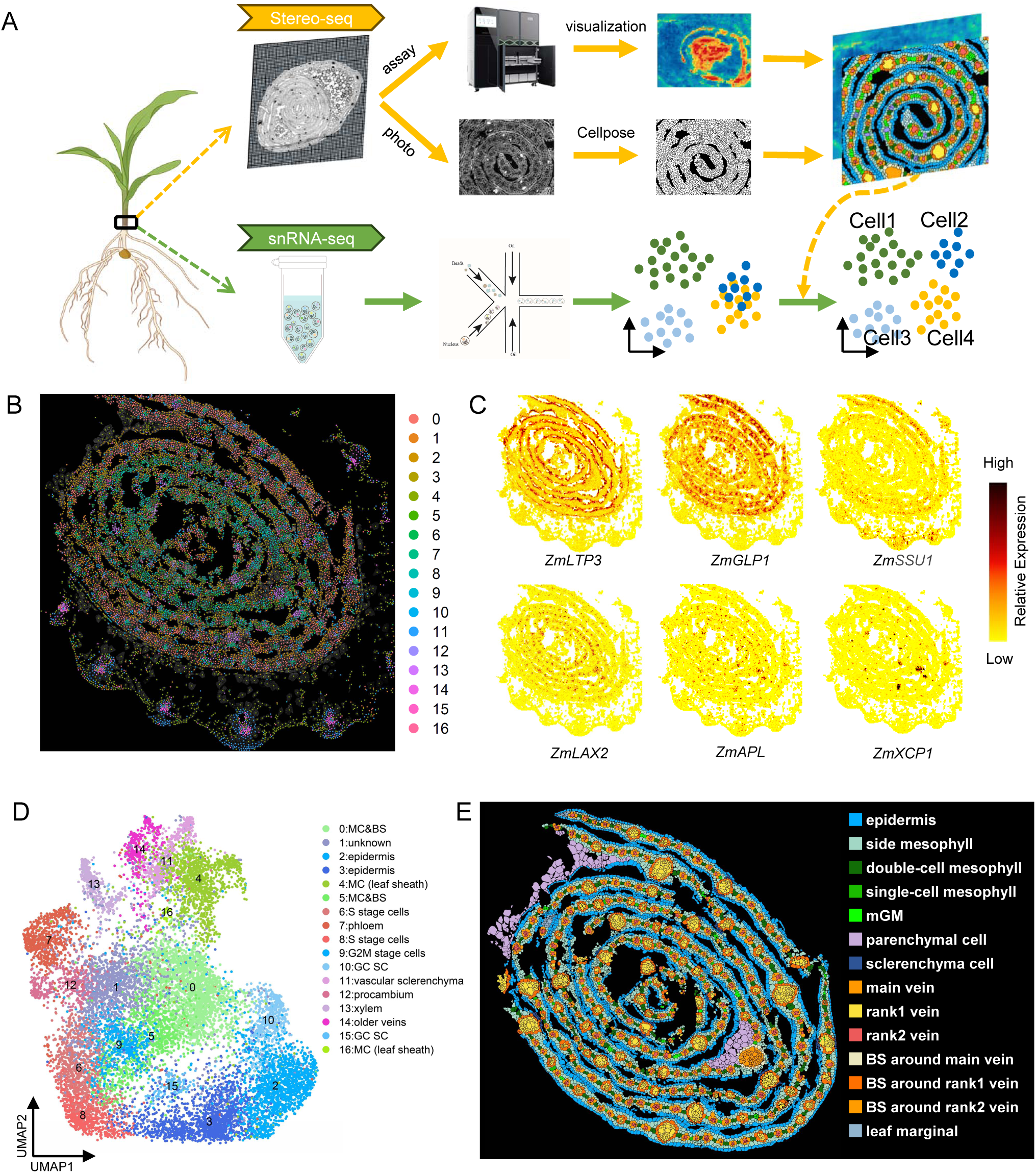
Spatial transcriptome of maize leaf primordiumx. (A) Workflow of Stereo-seq and snRNA-seq. The outlines of cell wall were recognized by Cellpose 2.0 and later be used to extract spatial single cell transcriptomes. Then the spatial marker genes were used to assist to distinguish the subtle differences between similar cell types of scRNA-seq data. (B) Spatial visualization of clusters on maize leaf primordium with outer leaf sheath. (C) Digital expression patterns of selected marker genes on Stereo-seq maps. Epidermis: *ZmLTP3*; mesophyll: *ZmGLP1*; procambium: *ZmLAX2*; phloem marker: *ZmAPL*; xylem marker: *ZmXCP1*; bundle sheath: *ZmSSU1*. (D) Annotated cell types of clusters identified on unsupervised clustering UMAP of maize leaf primordium with outer leaf sheath. (E) Recognized more (sub-)cell types based on the cell morphogenesis and spatial distribution on Stereo-seq maps of maize leaf primordium. mGM: median ground meristem.

The mesophyll cells (MC) and bundle sheath cells (BSC) are mixed in two clusters (C5 and C0) located in inner and outer primordium (Figure S1C). Given the morphologically distinct Kranz anatomy and the clear spatial arrangement of BSC and MC in an inner and outer circular pattern around the vein, respectively, we directly labelled and extracted their transcriptomes from Stereo-seq spatial data (Figure 1A and S2). Beside the cells of the leaf sheath, we recognized 14 (sub-)cell types in one section of the leaf primordium, including three BSC sub-cell types around three sub-cell types of veins (named main vein, rank1 vein and rank2 vein, respectively) and three sub-cell types of MC (side MC is next to epidermis, double-cell MC and single-cell MC are two and one cells of MC between veins) (Figure 1E, Figure S2). Then we identified differential expressed genes (DEGs) of all (sub-)cell types (Figure S3, Table S1). Based the fine classification of sub-cell type in the spatial transcriptome, we observed that single-cell MC preferentially expressed a higher number of genes associated with cell cycle processes, while double-cell MC and side MC exhibited higher expression levels of genes involved in photosynthesis (Figure S4, Tables S1-2). This result indicates that the three sub-cell types of MC are developmentally distinct in the primordium, which highlights the value of our single cell resolution spatial transcriptome.

### Identifying gradient expressed genes within the Kranz anatomy by a pseudo-Kranz strategy

In the leaf primordium section with spatial transcriptome data, three distinct stages of Kranz anatomy were identified: 544 Kranz anatomies with four BSC surrounding veins as 4BS-early stage, 813 Kranz anatomies with five unenlarged BSC as 5BS-middle stage, and 278 Kranz anatomies with enlarged BSC as 5BS-late stage (Figure 2A). ^31^ Since Kranz anatomy at the same developmental stages are distributed throughout leaves, it is challenging to derive the spatial relationships between genes, particularly low-expressed transcription factors (TFs). To better visualize and identify gene expression patterns within the Kranz anatomy, we superimposed the distributed Kranz anatomy of individual stages to create a unified “pseudo-Kranz”, with the center of each Kranz anatomy aligned at the origin of the coordinate system (Figure 2B, see Materials and Methods for further details). Unlike Kranz anatomy expression patterns in the spatial atlas, the expression of marker genes in the pseudo-Kranz is clearer and more vivid (Figure 2C; Figure S5).

**Figure 2.**
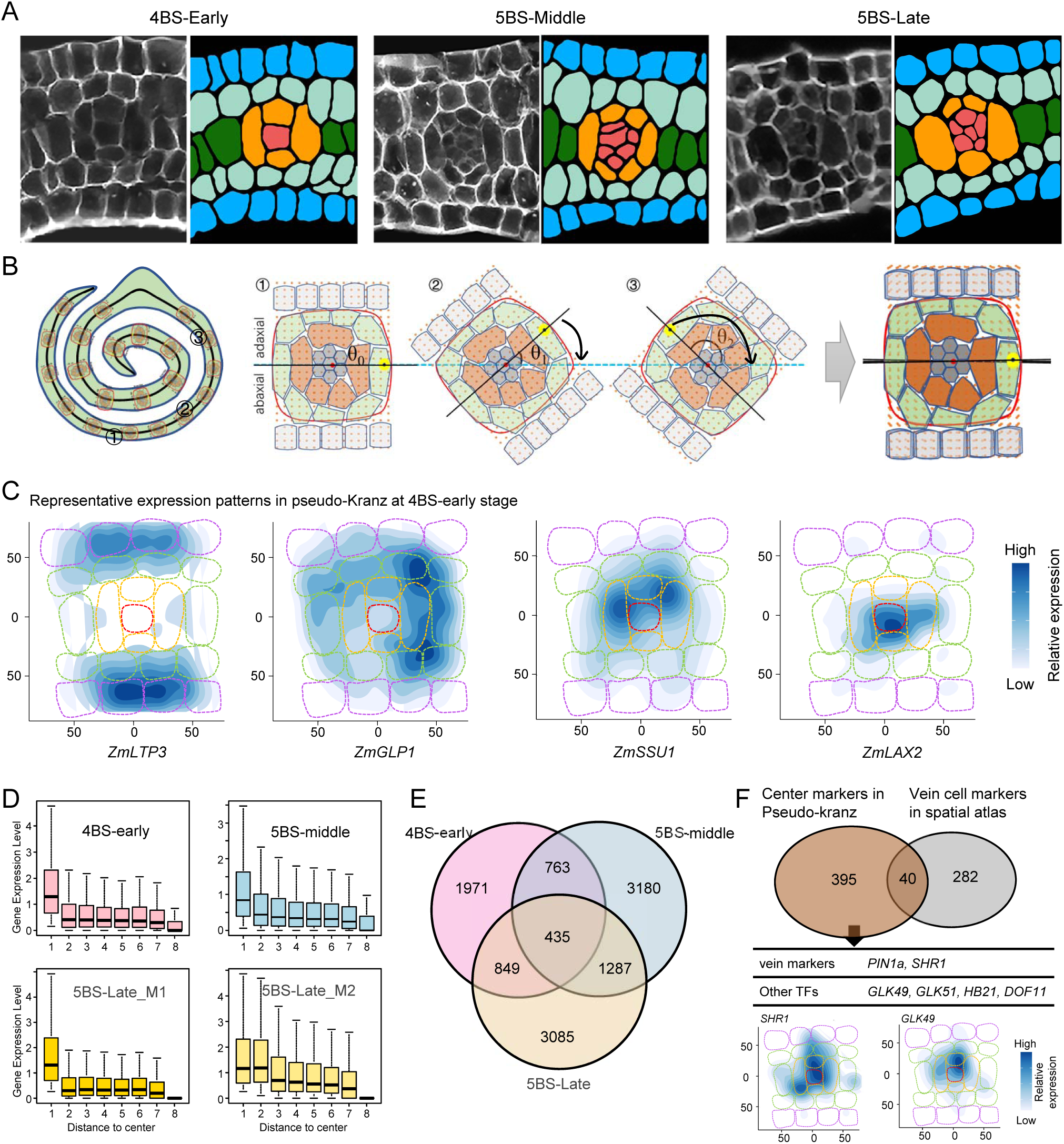
Integration of distributed Kranz anatomy into unified pseudo-Kranz. (A) Representative picture of three stages of Kranz anatomy in maize leaf primordium. Left: Fluorescence-stained images; Right: Recognized cell types with artificial color labels. (B) Workflow of integrating distributed Kranz anatomy at same stage into pseudo-Kranz. Left: Cartoon of spaced distribution of Kranz anatomy in leaf primordium; Middle: representative orientations of different Kranz anatomy which need to be rotated to horizontal direction; Right: overlap of different Kranz anatomy into pseudo-Kranz around original point. (C) Expression patterns of marker genes on pseudo-Kranz. Epidermis: *ZmLTP3*; mesophyll: *ZmGLP1*; bundle sheath: *ZmSSU1;* vein: *ZmLAX2*. (D) Modules with gradient decreasing patterns from center to periphery of pseudo-Kranz in three stages. One module with this expression pattern is detected either in 4BS-early or 5BS-middle stage and two modules are detected in 5BS-late stage. (E) Venn diagram of genes highly expressed in centers of pseudo-Kranz. (F) Representative new genes identified by pseudo-Kranz. Up: Venn diagram of markers identified in pseudo-Kranz and spatial atlas, respectively; Middle: representative new markers of pseudo-Kranz and bottom: two marker gene expression patterns on pseudo-Kranz.

The pseudo-Kranz analysis enables the identification of genes that exhibits a gradient of expression, progressing from the center to the periphery within the Kranz anatomical structure. To do this, we segmented the pseudo-Kranz into eight concentric rings with uniform width and calculated the average expression level of genes in each region. We then utilized weighted gene co-expression network analysis (WGCNA) to classify these genes into separate modules based on their expression patterns in the eight regions. We identified four modules of genes preferentially expressed in the center of pseudo-Kranz, encompassing 435 genes highly expressed at the center in all three stages (Figures 2D and 2E, Table S3). In comparison to DEGs of three vein cell types in the spatial atlas, a greater number of marker genes for veins were identified, for example, *ZmPIN1a*, *ZmSHR1*, *ZmGLK49*, *ZmHB21, ZmDOF11*, with the first two known for their role in regulating vascular bundle differentiation (Figures 2F).^9,32,33^

### Integrating spatial transcriptome with scRNA-seq atlas reveal the co-expression patterns of *SHR* and *SCR* genes in early BSC

To comprehensively understand the continuous development trajectory of Kranz anatomy across different stages, we complemented our Stereo-seq analysis with single-cell RNA sequencing (scRNA-seq) on identical sections (see details in materials and methods). After background RNA filtration with SoupX, 33,309 nuclei were obtained, and 30,548 genes were detected. These nuclei are clustered distinctly into 29 clusters, most of which are annotated with known marker genes (Figure 3A, Figure S6). In total, we found 8 clusters of epidermal pavement cells at different developmental stages (C2, C3, C10, C14, C8, C15, C23, C28) and two clusters of stomata cells (C22, stomatal precursor and C26, mature GC&SC). By integrating scRNA-seq data with Stereo-seq, we distinguished different vascular cell types: C16 and C27 represent xylem on the adaxial side of the vascular vessel; C7, C9, C11, C13, C19, C20 and C21 represent phloem on the abaxial side (Figure S7). C12 with the high expression levels of markers, such as *ZmLAX2*, *ZmMPa*, represents procambium cells; C18 with *ZmRS1*, *ZmLG3*, *ZmRss3* represents meristem; C1 with *ZmMAD2*, *ZmUBC19*, *ZmCYC23* represent G2M stage cells. C5 is assigned as MC expressing gene encoding *CA*, *PEPC* and *NADP-MDH* and C25 as BSC expressing genes encoding NADP-ME, SBPase and FBPase. C5 and C25 are more similar to MC and BSC in leaf sheath, respectively, rather than that of leaf primordium in spatial transcriptomics atlas (Figure S6).

**Figure 3:**
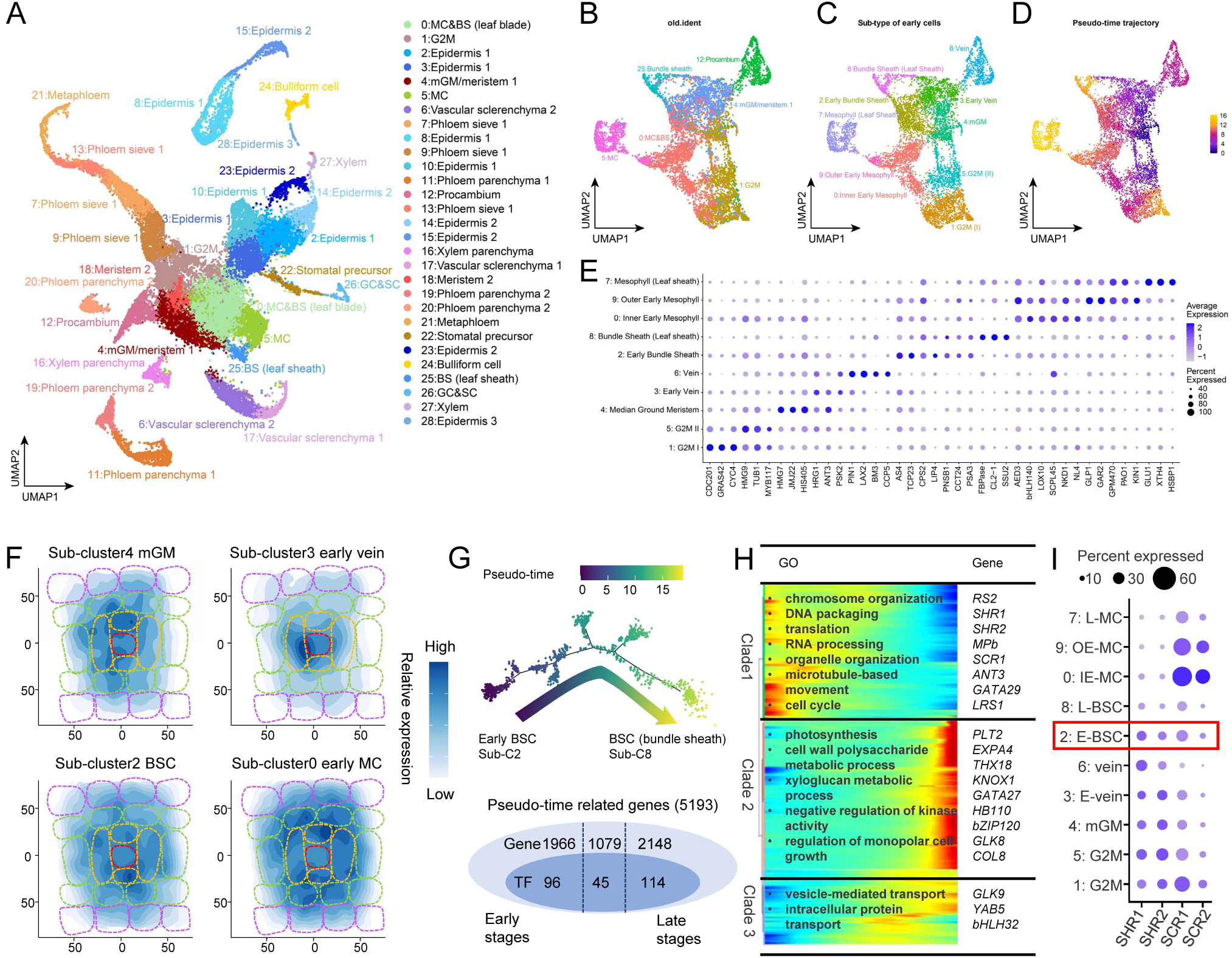
Integration of single-cell and spatial transcriptome attributes to identify early bundle sheath cell in maize leaf primordium. (A) The annotation of cell clusters on the UMAP of scRNA-seq atlas for maize leaf primordium with outer leaf sheath. (B) Supervised clustering of the subset cells at early developmental stages. The old annotation of the clusters of atlas are presented on the UMAP. (C) New annotation of sub-clusters on the UMAP. (D) Pseudo-time trajectory of subset cells displayed on the UMAP. (E) Dot plot of marker genes in the indicated cell types. The color key from light blue to dark blue indicates low to high gene expression levels of averaged scaled data. The dot size indicates the ratio of cells with expression compared to cells in that cluster. (F) The expression levels of DEGs identified in representative sub-clusters are summarized and visualized on 4BS-early pseudo-Kranz. (G) Pseudo-time analysis of early- and late-stage of BSC. Pseudo-time trajectory (up) and variable genes dynamically changed along the pseudo-time (pseudo-time related genes, bottom). (H) The expression heatmap of representative pseudo-time related genes and their enriched GO terms. (I) Dot plot of *SHR* and *SCR* genes in sub-clusters as showed in (E). IE-MC, inner early MC; OE-MC, outer early MC; L-MC, late MC. E-BSC, early BSC; L-BSC, late BSC; E-vein, early vein. The color key from light blue to dark blue indicates low to high gene expression levels of averaged scaled data. The dot size indicates the ratio of cells with expression compared to cells in that cluster.

Besides the MC and BSC clusters, we found that C0 also expressed relatively low levels of photosynthetic genes and relatively high levels of cell division related genes (Figure S6). We also found that C0 is more similar to the cells in primordium than that of leaf sheath (Figure S7). These results indicate that C0 possibly represents the mixed cells including BSC and MC at early developmental stages in primordium (Figure S7). To resolve the early MC and BSC, we subset four clusters at early developmental stages, including C0, C1, C4, C12, as well as MC of C5 and BSC of C25 by supervised clustering (see details in methods). We combined the 2000 variable genes identified by Seurat in these subset cells and DEGs identified in spatial transcriptome to conduct the dimensionality reduction of PCA analysis and then clustered these cells using top30 PCAs (Figure 3B). We assigned all 10 sub-clusters and distinguished MC and BSC at early developmental stages (Figures 3B-E). The sub-cluster 2 (sub-C2), a subset of cells comprising of cells within C4 and C0, was characterized by the high expression of marker genes preferentially expressed in the BSC identified in the spatial atlas (Figure 3E). We also found that sub-C2 is closely connected with sub-cluster of mature BSC in the pseudo-time trajectory analysis (Figure 3D) and have relatively high similarity with the cells around veins in the primordium (Figure S8). Furthermore, DEGs of sub-C2 were preferentially highly expressed in the region of BSC in pseudo-Kranz (Fig. 3F). The early MC were subdivided into Sub-C0 (inner early MC) and Sub-C9 (outer early MC), which are more similar to MC of very inner leaf primordium and outer leaf primordium, respectively (Figure S8).

The identification of these BSC at different developmental stages allowed us to study the developmental trajectory of early stage BSC (Figure 3G). There are total 5193 genes (including 255 TFs) expressed variably along the developmental pseudo-time trajectory (Figure 3G, Tables S4). As expected, BSC at early stage of pseudo-time expressed cell cycle-related genes, such as *ZmHIS2B5*, *ZmSMR1* and *ZmROA1* and enriched GO terms, such as “chromosome organization” and “DNA packaging” (Figure 3H, Tables S4-5). There are also a few of TFs, such as *ZmC78*, *ZmRS2*, *ZmGATA29* (*ZmGNC)*, preferentially highly expressed at the early stage of pseudo-time. At the late stages of pseudo-time, there was a progressive upregulation of photosystem genes and genes related to cell wall modification, such as *ZmPLT2*, *ZmXTH9*, *ZmEXPA4*, *ZmXT2* (Figure 3H, Table S4). Interestingly, we found that *ZmSHR1*, *ZmSHR2*, and *ZmSCR1* genes were both expressed at the early stage of pseudo-time in BSC (Figure 3H). Although *SHR* genes and *SCR* genes were predominantly expressed in early veins and MC, respectively, we observed their co-expression in early BSC (sub-C2) of scRNA-seq data (Figure 3I) and notably within confined single cells in spatial atlas (Figure S9). This suggests that, beyond their established roles in veins and MC^8–10,12,34^, *SHR* and *SCR* genes maybe also play important roles in establishing the identity of BSC in Kranz anatomy.

### Spatiotemporal co-expression network reveals the regulatory cascades underlying vein initiation and development

After we identified single cell transcriptome of most cell types of early-stage leaf, especially BSC at the early stage of Kranz anatomy, we conducted the co-expression network analysis of genes with Hotspot^35^ in these cells to mine key regulatory genes over leaf morphogenesis. Informative genes were filtered by gene autocorrelation and were clustered into 25 co-expression modules (Figure 4A, see details in methods). We identified specific modules associated with each cell type through mapping the expression patterns of each module in two ways: by mapping module scores of individual cells onto the UMAP of the scRNA-seq transcriptome and by displaying the total gene counts for each module on the pseudo-Kranz (Figure S10, Table S6). For instance, module 13 (M13, including *ZmSMXL3* and *ZmSHR1*, *ZmSHR2*) is associated with veins, which have high module scores in sub-C3 and sub-C4 and only highly expressed in the center of pseudo-Kranz. M7 (including marker genes, *ZmSSU2* and *G2*) is associated with BSC, which is featured with high module scores in the sub-C2 and preferentially highly expressed in the region of BSC in the pseudo-Kranz. M9 (including markers, such as *ZmGLP1* and *ZmCP29*) is associated with MC of Kranz anatomy (Figure 4B, Table S6).

**Figure 4.**
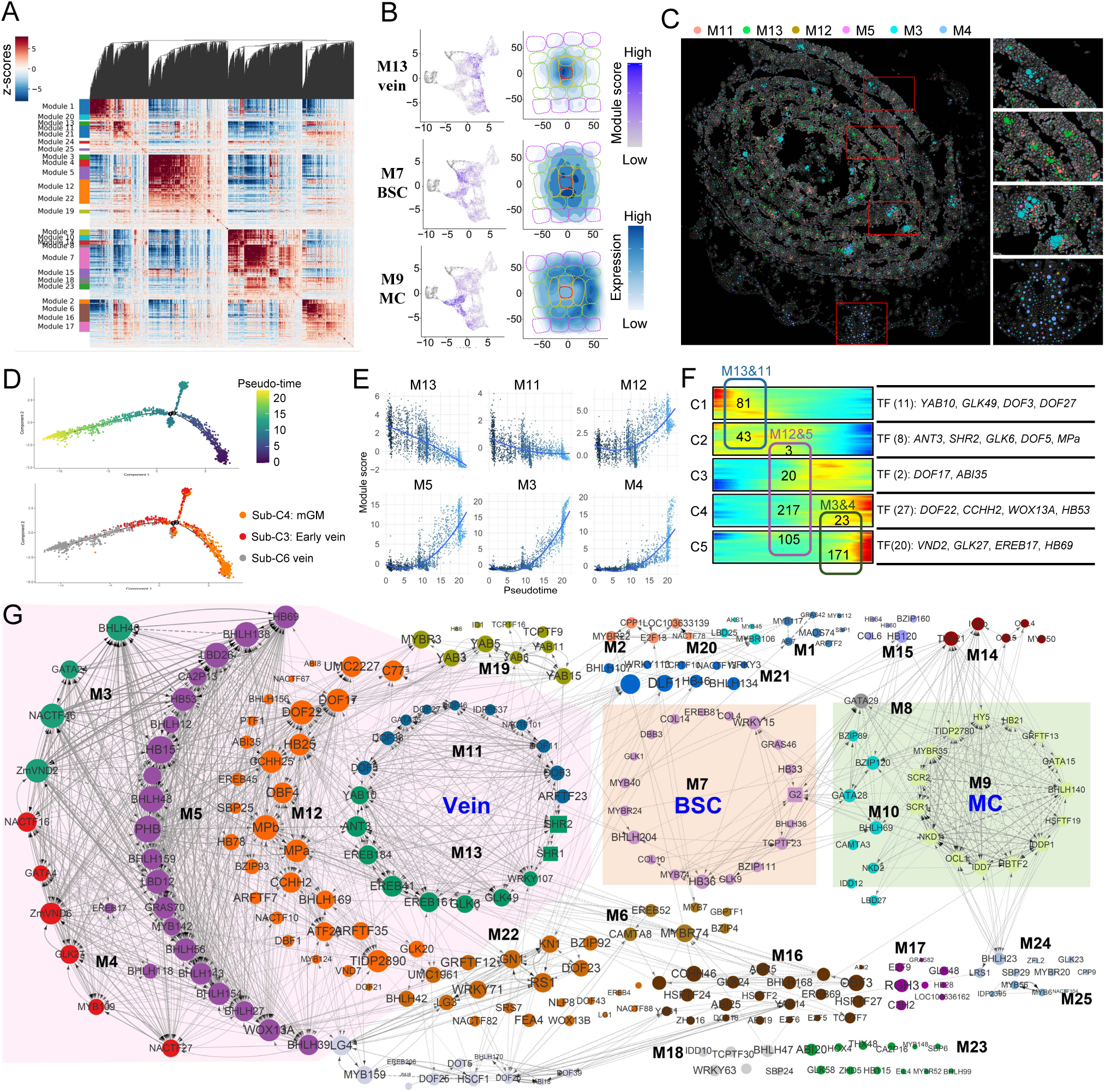
The regulatory network of maize leaf primordium. (A) Heatmap showing expression correlations between genes within 25 co-expression modules. (B) Total expression levels displayed on scRNA-seq UMAP and pseudo-Kranz. Left: Modules scores displayed on the UMAP of subset clusters, which represents the relative expression levels of each modules in each cells; Right: summarized expression levels of genes in modules showed on pseudo-Kranz. (C) The average expression of genes within vein-related modules on spatial atlas. The size of each spot represents the average scaled expression levels of genes within one module, with different colors indicating six distinct modules. Zoomed-in images on the right side showcase representative expression patterns of these modules. (D) Pseudo-time trajectory of sub-clusters of veins. Up: the pseudo-time of each cells in trajectory; Bottom: cell types of sub-clusters showed on the trajectory. (E) Module scores of each cell are showed along with the pseudo-time. (F) The variable marker genes dynamically along with the pseudo-time of vein were divided into five clades. Left: The numbers in box indicates the number of intersect genes with different vein-associated modules; Right: Representative TFs in the five clades of pseudo-time correlated genes. (G) Regulatory network in maize leaf primordium. Colors of nodes denote distinct modules. The size of nodes reflects the aggregate correlation score. Solid lines indicate predicted regulatory interactions between TFs, while dashed lines represent potential relationships without predicted binding sites. The “Zm” characters in front of the gene names have been omitted.

Interestingly, there were at least 6 modules (M3, M4, M5, M11, M12, M13) associated with veins, all of which have preferentially high expression levels in center of the pseudo-Kranz (Figure S10). To elucidate the spatial heterogeneity of gene expression for each module, we computed and graphed the mean normalized gene expression for each of the six vein-associated modules across the spatial atlas (Figure S11). By plotting the expression profiles of vein-associated modules across the spatial atlas, we found that M11 and M13 are predominantly higher expressed in small veins throughout the inner primordium, whereas M5 and M3 are mainly expressed in relatively mature lateral veins and midveins (Figure 4C). Additionally, M4 shows higher expression in more mature veins across the leaf sheath (Figure 4C). Consistent with the observed spatial heterogeneity, our analysis reveals that these modules are preferentially expressed in different sub-clusters of veins. For example, M3 and M4 have higher module scores in sub-C6 of vein, and M12 in sub-C6 and sub-C3, M13 in sub-C3, sub-C4 and sub-C5 (Figure S10).

We constructed the gene regulatory network (GRN) of veins within the leaf primordium by integrating co-expression relationships with potential TF-target interactions (see details in methods). The GRN clearly illustrates the regulatory cascades originating from TFs within modules M11 and M13, which subsequently influence modules M12 and M5, leading to downstream effects on modules M3 and M4. (Figure S12). It is noteworthy that key markers of vein initiation, including auxin-responsive factors (*ZmMPa* and *ZmMPb*) and auxin transporters (*ZmLAX2*, *ZmLAX3*, *ZmPIN1a* and *ZmPIN2*), are grouped in modules M12 and M5. This suggests that upstream regulators from M11 (such as *DOF* family genes) and M13 (such as *EREB* family genes, *ZmSHR1* and *ZmSHR2)* might influence vein initiation, thereby potentially controlling vein density. Additionally, there is a gene related to IAA transport, *BNLG1484*, which is the ortholog of *WAG1* and is potentially involved in establishing the auxin maximum during the initiation of veins and Kranz formation (Figure S12). This temporal regulatory network, spanning from vein initiation to development, provides a rich resource to support studying vein differentiation in maize (Figure S12).

To further dissect the regulatory cascades underlying vein development, we integrated our co-expression analysis with the construction of a developmental trajectory. This trajectory delineates the pseudo-time of developmental progression, tracing the path from median ground meristem (mGM, sub-C4) through early vein stages (sub-C3) to more mature veins (sub-C6) (Figure 4D). By mapping the module scores onto this trajectory for each cell, we observed distinct temporal expression patterns: M11 and M13 exhibits peak expression at early developmental stages, with expression levels subsequently declining as veins developed. Conversely, M12, M5, M3, and M4 display an inverse pattern, with expression increasing over the course of vein development (Figure 4E). The 4626 significantly variable genes along with pseudo-time can be divided into five clades by their expression pattern through pseudo-time (Table S8). By cross-referencing the genes from various vein-associated modules with those that exhibit pseudo-time-related expression patterns, we refined the pool of candidate genes involved in vein initiation and development (Figure 4F). Specifically, M11 and M13 overlap exclusively with genes expressed during the early pseudo-time stages, confined to clusters C1 and C2. In contrast, M12 and M5 show the overlap with genes expressed across stages C2, C3 and C4, while M3 and M4 aligned only with genes expressed during the later pseudo-time stages, specifically within clusters C4 and C5. These findings reinforce the existence of regulatory cascades spanning the six vein-associated modules and highlight potential master regulators governing the critical early phases of vein initiation (Figure 4F).

### Gene regulatory network identifies regulators of Kranz anatomy development in leaf primordium

For a systemic overview to identify new regulators involved in Kranz anatomy, we constructed a GRN of leaf primordium that includes all TFs within the co-expression modules (Figure 4G). As an application of this new transcriptomics atlas and cell specific GRN, in addition to a few of established markers, we identified many new regulators related to morphogenesis of Kranz anatomy. The *ZmSCR1* gene exhibits predominant expression in the MC of C_4_ plants (Figure S13), and its role in influencing the number of MC during Kranz anatomy development has been substantiated.^12^ Therefore, it is plausible to hypothesize that TFs co-expressed with *ZmSCR1* might act within the same regulatory pathway that regulates Kranz anatomy development. Among the most highly correlated TFs with *ZmSCR1* and/or *ZmSCR2* are *ZmNKD1*, *ZmIDD7*, *ZmIDDP1*, *ZmMYBR35*, *ZmHSFTF19*, *ZmBHLH23*, and *ZmNKD2* (also known as *IDD22*, Figure 4G). These candidate regulators were mainly expressed in early MC of leaf primordium but not in mature MC, suggesting that these TFs are very likely involved in morphogenesis and development of early-stage MC. It has been verified that *ZmSHR1* and *ZmSHR2* play crucial roles in the regulation of vein and MC development in Kranz anatomy.^7,9,17,34^ Within the GRN, a lot of TFs are predicted to directly regulate *ZmSHR1* and *ZmSHR2*. Notably, the auxin response factor *ZmARFTF23*, *ZmGLK49* and *YABBY* transcription factor *ZmYAB10* predominantly expressed within the mGM cluster (sub-C4), may affect the early expression of *ZmSHR1* and *ZmSHR2* during the initial stages of vein development in the mGM region (Figure 4G, Figure S12). Interestingly, we found that many TFs predicted to regulate *ZmSHR1* and *ZmSHR2*, such as*ZmGLK6*, *ZmYAB15* and *ZmDOF22*, exhibit dual expression in both vein and BSC, while some of the TFs in M7, the module particularly associated with BSC, were co-expressed in both BSC and MC (Figure 4G, Figure S13). The expression patterns indicate that there are possible regulatory cascades affecting the Kranz anatomy development from vein through BSC to MC.

### Comparing the scRNA-seq atlas between C_4_ and C_3_ leaves identifies the innovations for the developmental programs of Kranz anatomy

To identify new inventions during the evolution of C_4_ Kranz anatomy, we collected more scRNA-seq atlases for early developing leaves of C_4_ plants in grass, including *Sorghum bicolor* (hereafter called sorghum) and *Setaria viridis* (hereafter called Setaria) and one C_3_ plant of *Oryza sativa* (hereafter called rice) as control group. Similar to the annotations of maize scRNA-seq data, we identified most of the clusters of these three atlases through the expression patterns of known marker gene orthologs in each of these species (Figures S14-16, Table S9). In total, we obtained 24 (sub-)cell types in the four atlases, which include most of cell types in the sampled tissues (Figures 5A-D, Table S9). Even though each cell type occupied different proportion of cells in each species, we found highly conserved shared genes in each cell type, such as *PIN1a* and *LAX2* genes in procambium, *SUT1* in phloem parenchyma, *XCP1* in xylem (Figure D). Besides these shared marker genes in each cell type among these four species, we also identified many new marker genes for each cell type, which are conserved within four species (Table S10).

**Figure 5.**
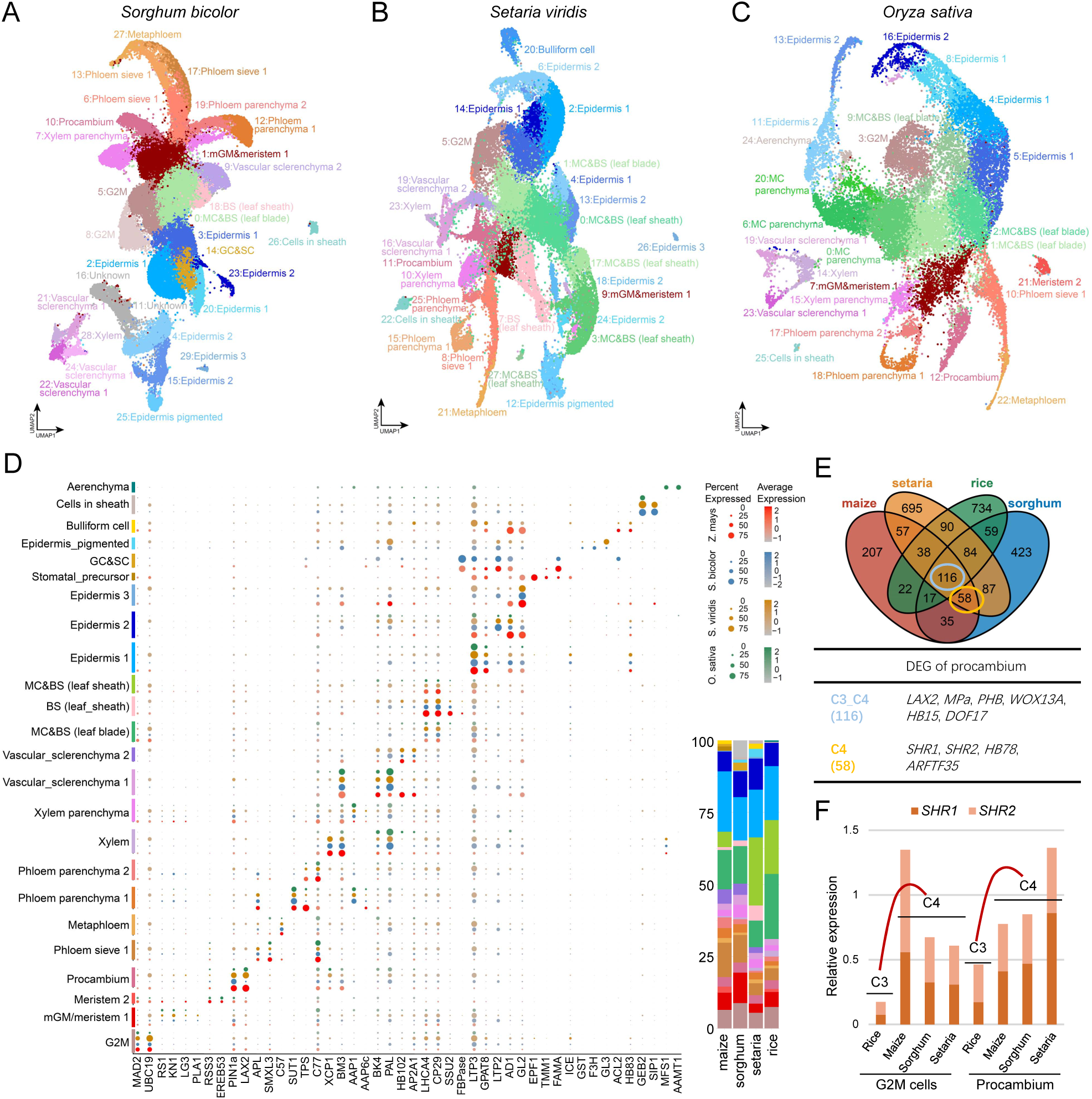
Comparison between scRNA-seq atlases of grass leaves. (A-C) The annotation of cell clusters on the UMAP of scRNA-seq atlases of leaf primordium with outer leaf sheath for sorghum (A), *Setaria* (B) and rice (C). (D) The expression patterns of conserved marker genes in common cell types identified in scRNA-seq atlases of the four grass species. The color key (red for maize, blue for sorghum, brown for Setaria and green for rice) from light to dark indicates low to high gene expression levels of average scaled data. The dot size indicates the ratio of cells with expression compared to cells in that cluster. The column at the bottom right shows the percentage of each cell type. (E) Comparison of DEGs within procambium between C_4_ and C_3_ species. Up: Venn diagrams showing the numbers of DEGs in different species; Bottom: C_4_ specific or C_4_ and C_3_ shared representative markers are listed in table. (F) The relative expression levels of *SHR* and *SCR* genes in G2M-stage cells and procambium identified in the four species.

The high level of overlap between the homologous genes in different cell types among the four species promoted us to further identify the new regulators involved in Kranz anatomy evolution. To do this, we first identified the conserved DEGs specifically present in each cell type of Kranz anatomy in C_4_ plants, then we compared these to that of corresponding cell type in rice to distinguish conserved genes among C_3_ and C_4_ plants or C_4_-specific markers. There were 106 DEGs shared by MC&BS among C_4_ and C_3_ plants (since here we could not differentiate MC and BSC in sorghum, Setaria, we used MC&BSC to represent the group including these two cell types), such as *NKD1*, *TCP23*, *G2*, *IDDP1*, *IDD7* and *ARFTF25*, that may perform the basic function during the development of MC and/or BSC (Figure S17, Table S10). In addition, we also found DEGs specifically identified in BSC&MC of C_4_ plant, such as *SCR1*, *SCR2*, *bHLH93*, and *IAA5* (Figure S17). The procambium is a key cell type whose initiation influences the vein density and development of Kranz anatomy. We found that procambium of three C_4_ species exhibited 174 common DEGs, with a large portion of 116 genes also present in rice procambium, while only a small subset of 58 genes was unique to the C_4_ plants (Figure 5E, Table S10). This result indicates a high degree of conservation in the developmental programs of procambium between C_4_ and C_3_ plants. We found not only many well-known marker genes associated with vein development, such as *LAX2*, *MPa*, *PHB*, *WOX13A*, but also many new markers, such as *HB124*, *DOF49*, *DOF17*, *bHLH48*, *NACTF46*, *LBD26*, *LBD12*, *RLD1* and *CCHH25*, which were shared among C3 and C4 plant. We also found some C_4_ specific marker genes of procambium, such as *SHR1*, *SHR2*, *HB78* and *ARFTF35* (Figure 5E).

Surprisingly, expression level of *SHR1* and *SHR2* genes in the procambium and G2M phase cells of C_4_ plants is significantly higher than that in C_3_ rice, just as the expression levels of *SCR* genes in MC&BS of C_4_ species are higher than that in C_3_ rice (Figure 5F and S17). Indeed, *SHR* genes are expressed in most cell types of vein in the three C_4_ plants, including phloem sieve, phloem parenchyma, xylem, and xylem parenchyma, procambium and even mGM, while rice *SHR* genes are only highly expressed in clusters of phloem sieve and xylem parenchyma in leaf and rarely expressed in procambium and other early-stage cells (Figure S18), indicating that *SHR* genes are initiated earlier and/or higher, resulting in a wider range of expression in C_4_ leaves than C_3_ leaves. Using public data of transcriptomics (https://www.plantrnadb.com/), we further found that the ratio of expression levels of maize *SHR* genes in leaf to that in root was higher than that of rice *SHR* genes (Figure S19).

The level of auxin in leaves is crucial for controlling vein patterning and leaf morphogenesis. We investigated key genes related to auxin, including biosynthesis, transport, and signaling (Tables S11-12). Contrary to expectations, we did not find significantly higher expression of biosynthesis genes or distinct expression patterns of signaling genes in C_4_ compared to C_3_ plants, which might be applied to explain the more veins in C_4_ than C_3_ plants (Figures S20-21). However, most IAA transport genes, including *PIN* genes responsible for auxin efflux and *LAX* genes for influx, showed preferential high expression in specific cell types (Figure S22). For instance, *PIN1a* gene were expressed in the procambium of both C_4_ and C_3_ plants, with relatively higher expression in C_4_ plants. In contrast, *PIN1b/c* genes were more expressed in C_3_ rice procambium. Notably, *PIN8* gene expression in C_4_ plant procambium was significantly lower compared to C_3_ rice, potentially influencing auxin levels and vein initiation in C_4_ plants (Figure S22).

### The developmental programs of Kranz anatomy are mostly recruited from C_3_ leaves with modifications

The GRN of early-stage leaves in the other three species were constructed using the same method as in maize. We then established a GRN for the developing leaves by comparing the conserved network in the three C_4_ plants with that of corresponding cells in the C_3_ rice (see details in methods). In this network, all known TFs involved in the development of Kranz anatomy are included. For instance, *DOT5*, also known as *TML1*, which was predicted to directly regulate *SHR1* and *SHR2* in this network (Figure 6A), was recently shown to specify small vein type in C_4_ and C_3_ grass leaves. *NKD1*, one of *IDD* family genes, which was predicted to directly regulate *SCR1* in this network, modulates *ZmSCR1* expression and impacts the MC number between veins in maize.^14^ This network also included additional *IDD* family genes that were predicted to directly regulate *SCR1* (Figure S23). Here, we verified the functions of other two *IDD* genes in the morphogenesis of Kranz anatomy in maize. We observed that the MC number between veins were significantly decreased in the *idd7* and *iddp1* mutants (Figure 6B, 6C and S24), comparable to those of *scr1* mutants.^8,12^ We then found that the expression levels of *SCR1* gene in both mutants of *idd7* and *iddp1* were lower compared with wild type control (Figure 6D), which suggests that *IDD7* and *IDDP1* participated in the development of Kranz anatomy, very likely through regulating expression levels of *SCR1* gene.

**Figure 6.**
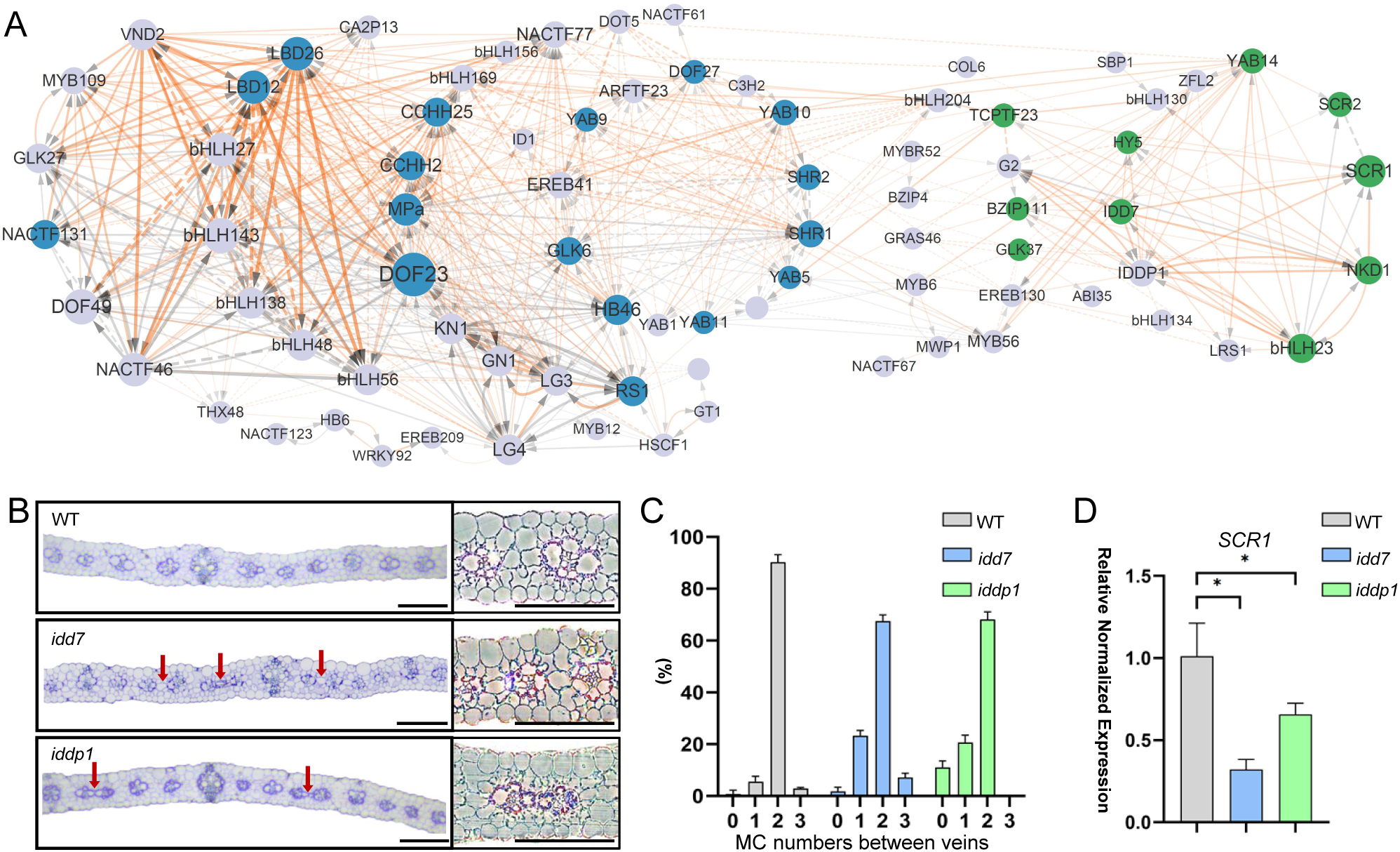
conserved gene regulatory network of Kranz anatomy. (A) Conserved co-expression network in grass species. The size of nodes reflects the mean correlation scores of four species. Orange lines represent specific conserved co-expression relationship in C_4_ species and gray lines represent the conserved co-expression relationship both in C_4_ and C_3_. Solid lines indicate predicted regulatory interactions between TFs, while dashed lines represent potential relationships without predicted binding sites. Blue nodes represents *SHR* genes and their TFs, while green nodes represents *SCR* genes and their TFs. (B) The transverse section of maize wild-type (WT, B73), *idd7* EMS mutant and *iddp1* EMS mutant. Red arrows indicate the loss of mesophyll cells or fusion of lateral veins in EMS mutants. Scale bar: 100μm. (C) Histograms summarizing the mean number of mesophyll cells separating veins in WT maize, *idd7* EMS mutants and *iddp1* EMS mutants, each for three replicates (N=3). (D) RT-PCR quantification of the expression of *SCR1* and *SCR2* in WT maize, *idd7* EMS mutants and *iddp1* EMS mutants. (*: P<0.05, two-tail Student’s t-test)

Analysis of this GRN shows that many conserved co-expression relationships of TFs in veins among three C_4_ plant were also found in C_3_ rice, which suggests a high level of conservation for the regulatory network for vein development between C_3_ and C_4_ (Figure 6A). As shown in the network, the relationship among many key hubs, such as *HB46*, *GLK6*, *EREB41*, *DOF23*, *MPa*, *BHLH156*/*48*/*138*/*143/27*, *DOF49*, *NACTF46*, *NACTF131*, *GLK27*, are conserved between C_3_ and C_4_ plants (represented as the gray lines in Figure 6A). Of course, there are also some genes, such as *SHR1*, *SHR2*, *DOF27*, *YAB10*, *ARFTF23*, *YAB9*, *NACTF77*, *BHLH169*, *CCHH25*, *CCHH2* and *DOT5*, which establish C_4_ specific co-expression relationships between each other gene and hence may play an important role in Kranz anatomy (represented as the orange lines in Figure 6A). These recruited regulators in the M11 and M13 include *YAB5*, *YAB10*, *YAB11*, *YAB9*, and *EREB41*, which are associated with the initiation and early development of veins (Figure 6A). We also identified the recruited regulatory relationships associated with MC in C_4_ leaves, such as those among *SCR1*, *SCR2*, *IDD7*, *NKD1*, and *IDDP1* (Figure 6A). By comparison, we identified only a limited number of novel regulatory relationships in BSC (Figure 6A), compared to either MC or vein development.

## Discussion

This study generated comprehensive spatial and scRNA-seq atlases for the young leaves of four grass species, including three C_4_ plants and one C_3_ plant as a control. Using these atlases, we successfully identified the transcriptomes of nearly all cell types present in the young leaves. The shared preferentially expressed orthologs suggests a high level of conservation of core functional pathways within the same cell types across grass species (Figures 5A-C, S17). These comprehensive atlases and the derived genetic regulatory networks combined with the pseudo-Kranz anatomy offer an opportunity to study the regulatory mechanisms associated with the development of different tissues or cells in a leaf.

Here we use the development of bundle sheath cell to further illustrate how these new data set allow us to identify regulatory mechanisms controlling differentiation of different cell type. BSC represents a key feature of C_4_ Kranz anatomy. In this study, by profiling the single-cell-resolution spatial transcriptomes, we can identify the subtle differences between early-stage BSCs and MCs, which were not distinguishable through unsupervised clustering of scRNA-seq data. The early stage of BSC shows characteristic expression of genes that are also preferentially expressed either in veins, such as *ZmSHR1*, *ZmSHR2*, *ZmMPb*, and *ZmANT3*, or in MC, like *ZmSCR1* and *ZmGATA29* (Figures 3H and S13), suggesting that early-stage BSCs share many regulatory relationships with both veins and MC. In addition to *ZmMPb* (Figure 3H), there are also many auxin-responsive factors that are differentially expressed at either the early- or late-stages of BSC, including 8 genes at the early stage and 18 genes at the late stage (Table S4), which indicates auxin plays a key role throughout the entire developmental trajectory of BSC. We further observed many *COL* family genes were expressed in late pseudo-time stages of BSC, such as *ZmCOL2*, *ZmCOL7*, *ZmCOL8*, *ZmCOL11* (Table S4). Therefore, *ZmCOL* genes might also be a major regulator of BSC development, possibly for chlorophyll accumulation, in addition to their other commonly known roles in flowering time control, responses to light, and auxin homeostasis in leaves.^36–38^

This pseudo-time series and derived GRNs allow us to construct a complete developmental trajectory controlling Kranz anatomy. The development of Kranz anatomy involves three cascades of events: initiation and development of veins, the differentiation of cell types, and cell specific development of photosynthetic machineries.^4^ The different cell types show drastically different levels of developmental complexity. For example, development of vein involves 6 modules, which represent a cascade of developmental changes (Figure 4G and S12). Several well-known marker genes related to initiation and early development of vein, such as *SHR1*, *SHR2*, and *ANT3*,^7,9,21,39^ are present along with a number of novel TFs, which include six *DOF* genes, three *EREB* family genes, two *GLK* genes and *WRKY107* (Figure 4G). Different regulators participated in middle- or late-stages of vein development as well (Figure 4G), e.g. *LBD* genes and *bHLH* family genes are enriched in M5, while *NAC* family genes are enriched in the late-stages of vein development. One notable phenomenon in the genetic regulation over vein development is that most of the involved genes have far more edges linking to other regulators, suggesting the high complexity of genetic regulation over vein initiation, compared to BSC or MC development. The developmental programs controlling BSC mainly involve M7, in which *G2* and *ZmTCP23* have the most linkages to other regulators. The developmental programs over MC involve M9, which includes many known regulators e.g. *ZmSCR1*, *ZmSCR2*, but also many new members, such as members of the *IDD* gene family, *HY5*, etc (Figure 4G). Here, we acknowledge that the well-known *SHR-SCR* regulatory module, which plays a crucial role in root development,^40^ also plays crucial roles in the development of Kranz anatomy, which is also supported by genetic evidences.^7–10,12,14,17,34^ It is worth emphasizing that these members identified to be involved in Kranz anatomy development need particular spatial and temporal combination to ultimately define the formation of Kranz anatomy. These identified modules and their members provide the first step for us to reconstruct the detailed genetic mechanisms controlling Kranz anatomy.

One of the major research goals of studying Kranz anatomy development is to identify the key evolutionary innovations promoting development of Kranz anatomy, which can be the most promising targets to engineer for the future purpose of creating a Kranz anatomy in C_3_ leaves. We tackled this challenge through a comparative analysis of genetic regulatory networks between C_3_ and C_4_ leaves (Figure 6A). Though BSC is commonly regarded as a special adaption in C_4_ photosynthesis, BSC in C_4_ leaves only contains a relatively few of new regulatory relationship in early developmental stage (Figure 6A). In contrast, MC in C_4_ leaves gained large number of new regulatory relationships. In particular, there are a large number of new regulatory connections developed between *SCR1* and its upstream regulators, including *IDD7*, *IDDP1*, and *NKD1* (also a member of IDD gene family), and surprisingly *HY5* (Figure 6A). Interestingly, these *IDD* family genes also confer a direct regulation over the expression of *G2*, an important regulator of the BSC development as well (Figure 6A).^5,6^ Possibly even more surprisingly, the vein development in C_4_ leaves gained the highest number of new regulatory relationships (Figure 6A), suggesting the primary importance of vein development in Kranz anatomy differentiation and should be highly researched to support future C_4_ engineering. Interestingly, in the modules controlling vein initiation, a large number of new regulatory connections were established between *SHR1*, *SHR2* and their regulators, including a number of *YABBY* family members, *EREB41*, *GLK6* etc. Considering that both *YABBY* family genes and *EREB* family genes are also closely related to drought responses of plants^13,41–45^ and these genes were upregulated in drought stress treatment (data from NCBI project number PRJNA421180 collected in transcriptome database http://ipf.sustech.edu.cn/pub/zmrna/),^46^ the genetic programs controlling adaption of C_3_ leaves to drought might be recruited to increase vein density during C_4_ evolution.^47,48^

Further examination of those expressional innovations in C_4_ leaves reveals mechanism controlling the evolution of Kranz anatomy. Compared to C_3_ leaves, *SHR* genes in C_4_ leaves are expressed at higher levels and across a broader range of cell types within the veins; the expression of *SCR1* increased in MC as well (Figures 5F and S18). Overexpression of *SHR* genes in both C_3_ and C_4_ plants typically result in an increase in the number of MC between veins and a decrease in the vein-to-MC ratio.^7,9,34^ The mutant *scr1* showed decreased number of MC between veins.^8,10,12,14^ Therefore, in C_4_ leaves, the changed expression of both *SHR1* and *SCR1* should have contributed to increased cell division to generate more MC between veins during leaf development. The decreased number of MC between veins during C_4_ plant evolution therefore can only be attributed to the increased initiation of procambium cells and hence minor veins. Auxin has been suggested as a major player in controlling the increased vein density in C_4_ leaves.^34^ However, in C_4_ grasses studied here, we did not notice increased expression of genes related to auxin synthesis (Figure S20). In contrast, a few genes related to auxin transport, e.g. *PIN1a*, *PIN8*, showed altered expression, which may contribute to elevated auxin levels in the ground tissues farther from the veins. Considering that *pin8* mutant cooperated with *pin6* increased vein branch points per area and *PIN8* overexpression exhibited simpler vein networks in *A*. *thaliana*,^26^ the decreased *PIN8* expression may be one of the key innovations to increase the vein numbers in C_4_ plants. These altered expression of either *SHR1* and *SHR2*, *SCR1*, and even auxin transport proteins are caused by the newly acquired regulatory relationship as discussed earlier, such as *DOT5* which possibly regulates *SHR1* and *SHR2* and three *IDD* genes—*NKD1*, *IDDP1*, and *IDD7*—which have been confirmed to regulate *SCR1* (Figure 6A).^14,49^ All these showcase a common theme of complex trait evolution, i.e. complex traits evolved through effective reutilization of existing mechanisms with changed expression patterns of key players. Such recruitment of existing regulatory mechanisms during the evolution of Kranz anatomy might underlie the repeated emergence of C_4_ photosynthesis during history.

### Limitations of the study

By comparing spatiotemporal scRNA-seq data in three C_4_ plant with C_3_ rice, we constructed a conserved gene regulatory network for early developing leaves. This enabled us to identify most of the known regulators for Kranz anatomy development, as well as numerous new TFs. Although we have validated two *IDD* family TFs affecting the number of MC between veins in Kranz anatomy, there are additional key hub genes in the network that need to be further verified for the evolutionary and developmental program of Kranz anatomy. In this work, we focused on the regulatory mechanism controlling the morphogenesis of Kranz anatomy. For fully elucidating the Evo-devo of C_4_ photosynthesis and engineering C_4_ system in C_3_ crops, it is also essential to reveal the regulatory mechanisms of establishing the C_4_ metabolism in mature Kranz anatomy.

## Data and code availability

The raw sequence data generated by Stereo-seq and scRNA-seq have been deposited in

CNGB Nucleotide Sequence Archive (project accession code: CNP0005465). To facilitate the application of this data source in plant biology research, we deposited the analyzed spatial data in Spatial Transcript Omics DataBase (STOmics DB) (ID STT0000093, https://db.cngb.org/stomics/project/STT0000093_5850b1c5/reviewer_link/).

All codes supporting the current study have been deposited in GitHub (https://github.com/jianzhaomichaelyang/STomics-C4Kranz).

Additional information required to reanalyze the data reported in this paper is available from the lead contact upon request.

## Acknowledgements

We thank Prof. Qihua Ling and Prof. Jiawei Wang from CEMPS for giving advice. We thank Xiaoyuan Wu from IBCAS for seeds of sorghum BTx623. We thank Bohan Zhang for help in Stereo-seq experiments; Fengqin Dong from IBCAS for help in semi-thin sections; Dr. Hui Zhang, Fuyu Li, and Feng Han, Yuhui Huang, Dr. Qingfeng Song and Linxiong Mao from CEMPS for help in data analysis and experiments; we thank Prof. Yongrui Wu, Dr.Jiechen Wang, Dr. Luhuan Ye, Lihao Chen, and Rui Yang for help in maize field planting; we thank Dr. Lina Shi, Xinyu Liu, and Fengping Ni for help in fund management. This study was supported by National Key R&D Program of China (No.2023ZD04073 to H.L.), Strategic Priority Research Program of the Chinese Academy of Sciences (grant number: XDB0630100 to XG.Z.), the general program of the National Science Foundation of China (31870214 to XG.Z.), the National Key Research and Development Program of China (2020YFA0907603 to XG.Z.) and the Shenzhen Science and Technology Program [KQTD20230301092839007 to C.T.] and the Science and Technology Major Special Project of Shenzhen (KJZD20230923114607016 to H.L.).

## Author contributions

X.G. Zhu and X. Xu conceived the idea; X.G. Zhu, X. Xu, Y. Gu and T. Wei supervised the work; X.G. Zhu, H. Liu, C.Y. Zhao and J.Z. Yang designed the experiments. H.Y. Chen, J.Z. Yang and C.Y. Zhao performed the Stereo-seq experiments with the help from W.W. Shao and L.C. Chen; J.Z. Yang, H.Y. Chen, H. Su, J. Yi, Y.T. Cheng and Y. Wang performed cell identification of Stereo-seq data. A.T Ni and C.Y. Zhao performed the scRNA-seq experiments. C.Y. Zhao, J.Z Yang, A.T. Ni, H.Y. Chen, H. Su and L.Y. Zhong performed data analysis. D.M. Fang, H.J. Li, H.X. Sun, Y.Q. Bai, S.S. Ma and X.Y. Tu provided essential advice for analysis. J.Z. Yang, Y.T. Cheng and X.Y. Li identified maize EMS mutants. J.Z Yang, Y.T. Cheng, H.R. Sun and Y. Dong performed resin section and microscopy imaging. C.Y. Zhao, J.Z. Yang, A.T. Ni and H.Y. Chen prepared the figures. C.Y. Zhao, J.Z. Yang, A.T. Ni and H.Y. Chen prepared the manuscript with help from all authors. X.G. Zhu, X. Xu, Y. Gu, T. Wei, H. Liu, P. Wang, G.Y. Zhang and X. Guo provided valuable advice and reviewed the manuscript. X.G. Zhu, X. Xu, Y. Gu and T. Wei finalized the manuscript. All other authors contributed to the work. All authors approved the manuscript for submission.

## Supplemental information

Figure S1. Spatial transcriptome of maize leaf primordium

Figure S2. Representative picture of cell type recognized in spatial transcriptome

Figure S3. Expression patterns of marker genes identified in each cell type on Stereo-seq maps

Figure S4. Expression patterns of photosystem related genes in three sub-types of mesophyll cell

Figure S5. Expression patterns of marker genes of four cell types on three stages of pseudo-Kranz

Figure S6. Dot plot of marker genes in the indicated cell types of maize leaf atlas

Figure S7. The integration of clusters of snRNA-seq atlas with cells of spatial transcriptomes of maize

Figure S8. The integration of sub-clusters of snRNA data with cells of spatial transcriptomes

Figure S9. co-expression patterns of SHR and SCR genes in spatial transcriptomes

Figure S10. Expression patterns of module genes in snRNA-seq and pseudo-Kranz

Figure S11. Expression patterns of individual vein-associated module genes spatial atlas

Figure S12. Regulatory co-expression network of vein related module genes

Figure S13. The expression patterns of candidate regulators of SHR, SCR and G2 genes

Figure S14. Dot plot of marker genes in the indicated cell types of sorghum leaf atlas

Figure S15. Dot plot of marker genes in the indicated cell types of Setaria leaf atlas

Figure S16. Dot plot of marker genes in the indicated cell types of rice leaf atlas

Figure S17. Conserved DEGs in BSC and/or MC among C4 and C3 plants

Figure S18. The expression patterns of selected key genes in different cell types of C4 and C3 plants

Figure S19. The expression levels of SHR genes in root and leaf of maize and rice

Figure S20. The expression levels of auxin biosynthesis genes in C4 and C3 species

Figure S21. The expression levels of auxin signal transduction genes in C4 and C3 species

Figure S22. The expression levels of auxin transport genes in C4 and C3 species

Figure S23. The relative positions of predicted binding sites of TFs co-expressed with SCR1

Figure S24. The mutation site identification of EMS mutants

Table S1. The differentially expressed genes identified in each cell type in spatial transcriptomes of maize early leaf

Table S2. The average expression of photosynthesis related genes in each cell type of spatial transcriptome of maize early leaf

Table S3. Spatial heterogeneity of gene expression in three stages of pseudo-Kranz

Table S4. Marker genes variable along with pseudotime of early BS (sub-cluster 2) and relative late BS (sub-cluster 8) in snRNA-seq transcriptome

Table S5. GO terms enriched in each cluster of marker genes variable along with pseudotime of early BS (sub-cluster 2) and relative late BS (sub-cluster 8) in snRNA-seq transcriptome

Table S6. The gene modules of snRNA-seq transcriptome of early maize leaf calculated by hotspot

Table S7. The GO terms enriched for genes in each module of snRNA-seq transcriptome of early maize leaf calculated by hotspot

Table S8. Marker genes variable along with pseudotime of veins related clusters (sub-C4, sub-C3 and sub-C6) in snRNA-seq transcriptome

Table S9. Marker genes were used to annotate the cell types in scRNA-seq atlas of four species

Table S10. The conserved DEGs of each cell type in C_4_ plants

Table S11. The orthologs of IAA related genes in four species

Table S12. Mean expression of IAA related genes in the clusters of atlas in four species

Table S13. Annotation and gene names of maize genes

Table S14. Orthologs of maize genes in Sorghum bicolor, Setaria viridis and Oryza sativa subsp. Japonica

Table S15. Primers list for EMS mutants genotyping and RT-PCR quantification

Table S16. Datasheet of RT-PCR quantification

Table S17. Datasheet of counting number of mesophyll cells separating veins

## EXPERIMENTAL MODEL AND STUDY PARTICIPANT DETAILS

### Plants

In this study, we utilized four species: three C_4_ plant *Zea mays* - B73; *Sorghum bicolor*-BTx623; *Setaria viridis* - A10) and one C_3_ plant (rice, *Oryza sativa japonica*). Plants were cultured in greenhouse of China National Gene Bank (CNGB), Shenzhen, with an average temperature of 28℃ and 55% humidity was kept. The average light intensity was maintained at 500 μmol·m^-2^·s^-1^ for 16 hours daylight and 8 hours dark. Plant tissues were sampled when maize grew around 12 days, sorghum around 12 days, Setaria around 10 days, and rice around 15 days, reaching the vegetative stage with three leaves. Tissues, approximately 5 mm above from ground, comprised apical meristem plus leaf primordia from P1 to P6, were harvested for both Stereo-seq and scRNA-seq analyses. For maize (*Zea mays)*, the outmost leaf sheath was stripped off.

## METHOD DETAILS

### Stereo-seq

To spatially visualize transcriptomic expression within leaf tissue, the STOmics gene expression reagent kit was used, following the protocol described by the Stereo-seq chip manufacturer.^48^ The fresh tissue was immediately embedded in PVAL - OCT (PVAL by Shanghai M&G Inc., No. MF-7001, and OCT by SAKURA, No. 4583) and then frozen with liquid nitrogen-precooled isopentane. Cryosections, adhered to and fixed on the chip, were stained with Qubit™ ssDNA Reagent (SS, Invitrogen, No. Q10201) and Fluorescent Brightener 28 disodium (FB, Sigma, No. 910090), followed by fluorescence imaging. The tissue is then permeabilized based on the tissue-specific optimization protocols (BGI, No. 1000028492): 10 minutes for maize and sorghum, 30 minutes for Setaria viridis, and 6 minutes for rice. Subsequent procedures were carried out according to the STOmics Gene Expression Kit-S1 instructions (BGI, No. 1000028493 for library construction and sequencing via the MGI DNBSEQ platform in CNGB.

### Raw Stereo-seq data processing

Sequenced reads were generated using an MGI DNBSEQ-T10 sequencer and the expression profile matrices with coordinate on chips were produced according to the Stereo-seq method.^28^ The reads of CID and MID were used to label cells and locate coordinates, respectively. The remaining transcript reads were then aligned to genome references (maize, Zm-B73-REFERENCE-NAM-5.0.55 from maizeGDB; sorghum, *Sorghum bicolor* v3.1.1 from Phytozome; Setaria, *Setaria viridis* v2.1 from Phytozome; rice, *Oryza sativa* v7.0 released by MSU) to generate gene expression raw matrix.

### Image Registration and Cell Identification

The raw matrix was used to generate gene expression maps on chip, which includes expression counts of each gene on individual DNBs. The SS image was registered to the gene expression maps based on the track lines on chip with an average error less than 0.5 um, and then overlaid with the FB image to align the tissue contours. Cell segmentation and identification were conducted based on FB image using the Cellpose 2.0 segmentation algorithm.^50^ A custom model was trained using a Nvidia RTX3080Ti GPU based on the cyto2 initial model. After 12 iterations supervised by Human-in-the-Loop annotations, the model achieved a recognition accuracy exceeding 95%. The identified cell outlines were converted into grayscale bitmaps. These bitmaps were overlaid with the original microscopic images using Fiji ImageJ. Cells were manually classified based on anatomical structures and marked with different colors. Masks corresponding to different types of cells were exported along with their coordinates. A Python script was employed to extract single-cell stereo-seq transcriptomes based on image-defined boundaries for downstream analysis.

### Unsupervised clustering of Stereo-seq transcriptome and annotation

The MID counts of each gene captured by DNBs in cell segmentation were extracted based on the masks of easch tissue on chip and summarized to generate gene expression matrix for downstream analysis. Then the data of cell-gene matrix was processed in Suerat (v4.4.0).^51^ Cells with MID counts less than 10 were discarded and raw data of counts were normalized by SCTransform function. A KNN graph was constructed by FindNeighbors function with 30 dimensionalities of PCA and cluster analysis was performed by FindClusters function with resolution of 1. Clustering results were displayed by uniform manifold approximation and projection (UMAP) dimension reduction analysis. Established marker genes were then utilized to annotate the cell types of clusters. For more accurate cell annotation, the labeled parts of cells were directly recognized based on the relative positions in the section of leaves. These cell identities were then added into the metadata of clustering data in Seurat (v4.4.0).

### Integrate transcriptome of same stage Kranz anatomy into Pseudo-Kranz anatomy

The points of center and right-middle of each Kranz anatomy were labeled to establish the middle transverse line with symmetry on both adaxial and abaxial sides. Then rotated the Kranz anatomy to same horizontal line and to integrated same stage Kranz anatomy into pseudo-Kranz by converting the center points of each Kranz anatomy to origin of coordinates (0, 0). The expression patterns of individual genes or total of several genes are showed as contour lines by plotting the MID counts on the pseudo-Kranz.

### Preparation of nucleus suspension

The protocols were modified based on former works.^52^ Tissues were cut into small pieces in pre-cooled buffer B (0.4 M sucrose, 10 mM MgCl_2_, 25 mM Tris-HCl pH = 8.0, 1% RNase inhibitor, 0.1 mM PMSF, 1% Triton X-100, 0.1% DTT) and then incubated on ice for 10 minutes and filtered. with addition of 0.05% TritonX-100 for maize and 0.1% TritonX-100 for sorghum, Setaria and rice.

For maize, the nuclei were centrifuged for 10 minutes (1000g, 4°C, 5g/s) and resuspended in Buffer B. A 5-fold volume of Buffer F (1X PBS, 1% RNase inhibitor and 0.1% DTT (BGI, 01B022BM)) was added and centrifuged for 10 minutes (1000 g, 4°C, 5 g/s) to remove most organelles and other molecular impurities. For Sorghum, Setaria, and rice, large cell debris was centrifuged down to the bottom at 200g for 5 minutes (4°C), and the nuclei in the supernatant were centrifuged at 1500 g for 20 minutes (4°C) and resuspended in Buffer B again. Density gradient centrifugation (3000g, 4°C, 30 minutes) was performed using 30% percoll (Biosharp, BS909) and 80% percoll to remove organelle and nuclei included in the middle layer was washed by adding 5 times the volume of Buffer F. The precipitate was resuspended with cell resuspension solution to obtain the nuclei resuspension for downstream microfluidic experiments. 2µl cell nuclei resuspension and 8µl DAPI (Beyotime, C1005) solution was mixed well and counted under fluorescence microscope. Only intact nuclei were counted.

### scRNA-seq and obtaining gene expression matrices

After obtaining the nuclear suspensions, microfluidic experiment and library preparation were conducted according to the DNBelab C Series Single-Cell Library Prep Set (MGI, No. 1000021082) as previously described.^53^ Both library construction and sequencing were performed on the MGI DNBSEQ-T1&T5 platforms. The obtained raw data was first analyzed using single-cell DNBelab C4 technology (v2.0)^54^ of MGI to automatically obtain two gene matrices: RawMatrix including reads of all DNBs and FilterMatrix including reads of filtered DNBs. The genome references were same as that used at analyses of Stereo-seq. The RawMatrix and FilterMatrix were further processed using the SoupX algorithm to remove background contamination to generate final expression matrix for downstream analysis.^55^

### Quality control and unsupervised clustering of scRNA-seq data

Cell-gene expression matrices were processed and unsupervised clustering using Seurat (v4.4.0).^51^ For filtering high-quality data of cells, genes expressed in less than 3 cells and cells with less than 2000 in maize, 1000 in sorghum and 500 in Setaria and rice were removed. The data from different matrices of same species were merged and then normalized using the NormalizeData function with the “LogNormalize” method (scale.factor = 10000). Cell cycle phase were predicted Using the CellCycleScoring function in order to mitigate adverse impact on downstream analysis of clustering.^56^ PCA dimensionality reduction was performed and calculate top 30 principal components using RunPCA function with 3000 variable genes identified by FindVariableFeatures function. Batch effects were removed using RunHarmony function. Unsupervised clustering was conducted utilizing the FindNeighbors and FindClusters functions with a resolution parameter set to 1, and the results were visualized using Uniform Manifold Approximation and Projection (UMAP).

### Integrate scRNA-seq transcriptome with stereo-seq atlas

We integrated the scRNA-seq transcriptome with the stereo-seq atlas using the TransferData function in Seurat (v4.4.0).^51^ Prior to integration, both datasets were normalized using the SCTransform function. Anchors were calculated using the FindTransferAnchors function, and predictions.assay of integration were performed using TransferData function and added to the metadata of the spatial atlas. The correlation of clusters with cells from spatial data was plotted on the spatial atlas using the SpatialFeaturePlot function.

### Cell type annotation in scRNA-seq atlas

We firstly collected known markers that are preferentially expressed in specific cell types either in maize and rice or other species and obtained their orthologs in maize, sorghum, Setaria and rice. Most of cell types occurring in the four species were identified using known markers (Table S12) and were further confirmed by the results of integration of scRNA-seq and spatial transcriptome. For the few clusters without well-known markers, we primarily relied on the integration of scRNA-seq and spatial transcriptomics to infer cell types. Additionally, we considered the orthologs of DEGs that are specifically expressed in these clusters and are conserved across different species.

### Subset of single cells at early developmental stages and supervised clustering

The clusters of early stages of cells in scRNA-seq transcriptome were subset based on the annotation of the atlas and re-clustered as the method of clustering for atlas, except the parameter of resolution in FindClusters functions were set to 0.5. For supervised clustering to distinguish the early bundle sheath cells (BSC) of scRNA-seq data in maize, the PCA were conducted using RunPCA function with variable gene set that consist of 1, 478 DEGs identified in spatial transcriptome and 2, 000 variable genes identified by FindVariableFeatures function. The following steps of dimension reduction and clustering were performed as the method of clustering for atlas.

### Identifying conserved DEGs specific in C_4_ spieces or in both C_3_ and C_4_ spieces

Firstly, we identified differentially expressed genes (DEGs) for each cluster across using the FindMarkers function in Seurat (v4.4.0).^51^ Subsequently, we derived the conserved marker gene sets for each cell type across the three C_4_ species by intersecting the orthologous genes among the DEGs identified in this cell type. We then identified cell type markers that are specific to C_4_ species or shared between C_3_ and C_4_ plants by comparing the conserved marker gene sets of each cell type in C_4_ species with those of C_3_ rice.

### Developmental trajectory analysis

The developmental trajectory analyses were conducted by employing implemented function in Monocle 2 (v2.18.0)^57^ or Monocle 3 (v1.0.0)^58^ for different samples. For the entire cell population of the early stage of leaf subset, Monocle3 was employed to delineate the developmental trajectories of different cell types. Meanwhile, for individual cell types, Monocle 2 was utilized to primarily compute the pseudo-time of each cell along its developmental path. The transcriptomes of the subset cells were analyzed using Seurat (v4.4.0) for dimensionality reduction and subsequent clustering. For Monocle 2, a matrix of gene expression counts was utilized to create a new CellDataSet object using the newCellDataSet function, with the expressionFamily set to negbinomial.size(). Subsequently, size factors and dispersions were estimated using the estimateSizeFactors and estimateDispersions functions, respectively. Genes for reducing data dimensionality and ordering cells were identified using the differentialGeneTest function, based on clusters established by Seurat (v4.4.0). The genes were filtered using the parameters q value < 0.01 and mean_expression >= 0.1. Cell order was conducted using orderCells with default parameters. Genes that Change as a Function of Pseudotime were found using differentialGeneTest function. For Monocle 3, the gene expression matrix was normalized using the preprocess_cds and align_cds functions. Dimensionality reduction and clustering were performed using the built-in functions of Monocle 3. The developmental trajectory was reconstructed with the learn_graph function, and cells were subsequently ordered in pseudo-time using the order_cells function.

### Construct co-expression regulatory network

To investigate the transcriptomic features of the bundle sheath and mesophyll cell arrangements in C_4_ plants, the Python package ‘Hotspot’ was employed for functional gene module analysis.^35^ Initially, single-cell data were filtered to retain the top 5000 most highly variable genes, along with genes of interest (some genes of interest may not be among the highly variable genes and thus need to be manually added to a gene list to prevent their exclusion). The ‘danb’ model was utilized, and modules were identified based on an FDR threshold of <0.00001 and genes of interest. The create_modules function was then used to recognize modules with a min_gene_threshold of 40 and an fdr_threshold of 0.05. Module scores were calculated using the module_scores function to measure the gene expression levels in each of cells. Subsequently, a co-expression network between different genes was constructed using autocorrelation z-scores and visualized in Cytoscape. The regulatory relationships of TFs with targets were predicted based on two databases of TF binding sites (JASPAR, https://jaspar.elixir.no/; CIS-BP, http://cisbp2.ccbr.utoronto.ca/index.php). Then the regulatory relationships of TF and target with predicted binding sites were added into the co-expression relationship network to construct GRN.

### EMS mutants genotyping

T3 generation seeds of B73 maize background, treated with EMS mutagenesis, were ordered from the mutant library available at http://maizeems.qlnu.edu.cn. After sowing, leaf tissue was collected from the seedlings on the fifth day for DNA extraction (TIANGEN DNAquick Plant System, No. DP321-03) followed by PCR-based genotyping (2×Taq Plus Master Mix II, Vazyme, No. P213-03) to identify mutant genotypes and select homozygous lines.

### RT-PCR quantification analysis

Plant tissues were sampled and ground after freezing in liquid nitrogen. RNA was extracted using the GeneJET Plant RNA Purification Kit (Thermo Fisher Scientific, No. K0802). The extracted RNA was reverse transcribed to generate cDNA using the TransScript® All-in-One First-Strand cDNA Synthesis SuperMix for qPCR kit (TransGen Biotech, No. AT341-01), with 1000 ng of cDNA synthesized per sample. Diluted cDNA samples were subjected to real-time quantitative PCR using the PerfectStartTM Fast Green qPCR SuperMix (TransGen Biotech, No. AQ611-04), with the HSP-96 well plate (Bio-Rad, No. HSP9601) and the CFX Connect Real-Time system (Bio-Rad). All quantifications were normalized to maize β-Tubulin1 (Zm00001eb000490) as an internal reference. Data analysis was performed using Bio-Rad CFX Maestro software (Version: 4.0.2325.0418). Each quantitative experiment was conducted with three biological replicates, each containing three technical replicates. Normalized expression analysis (ΔΔCq) was performed based on biological replicates, with significance determined using a t-test with a p-value < 0.05.

### Resin section and microscopy imaging

Plant tissues were sampled and immersed in FAA solution (a solution containing 50% ethanol, 5% acetic acid, and 3.7% formaldehyde in water) and subjected to vacuum infiltration at 0.4 MPa on ice to remove bubbles. Subsequently, the plant samples were dehydrated in a gradient ethanol series and embedded using the Technovit 7100 Embedding Kit (Heraeus Kulzer, No. HCECS000677). After solidification, 8 mm thick sections were cut using a Leica UC7 microtome, and the sections were floated onto adhesive-coated slides. Cell wall autofluorescence was observed using an Olympus fluorescence microscope, and images were captured.

---

**Supplementary figure 1.**
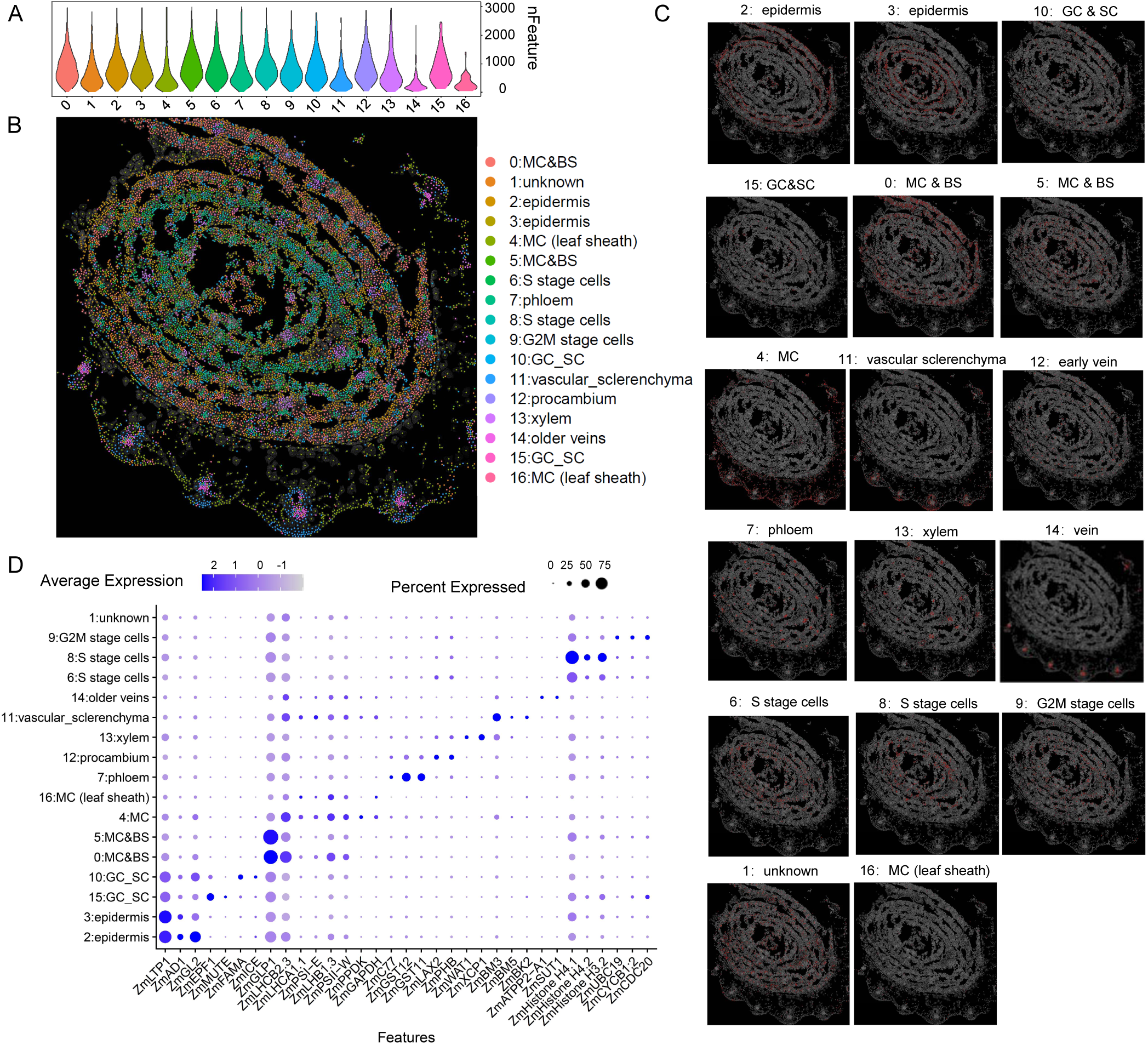
Spatial transcriptome of maize leaf primordium. (A) Violin plot showing the number of genes detected in each cell across different clusters in the spatial atlas. (B) Cell annotation on the spatial atlas of maize leaf primordium including the outer leaf sheath. (C) Cells were separately displayed on the Stereo-seq maps. (D) Dot plot of marker genes in the indicated cell types. The color key from light blue to dark blue indicates low to high gene expression levels of averaged scaled data. The dot size indicates the ratio of cells with expression compared to cells in that cluster.

**Supplementary figure 2.**
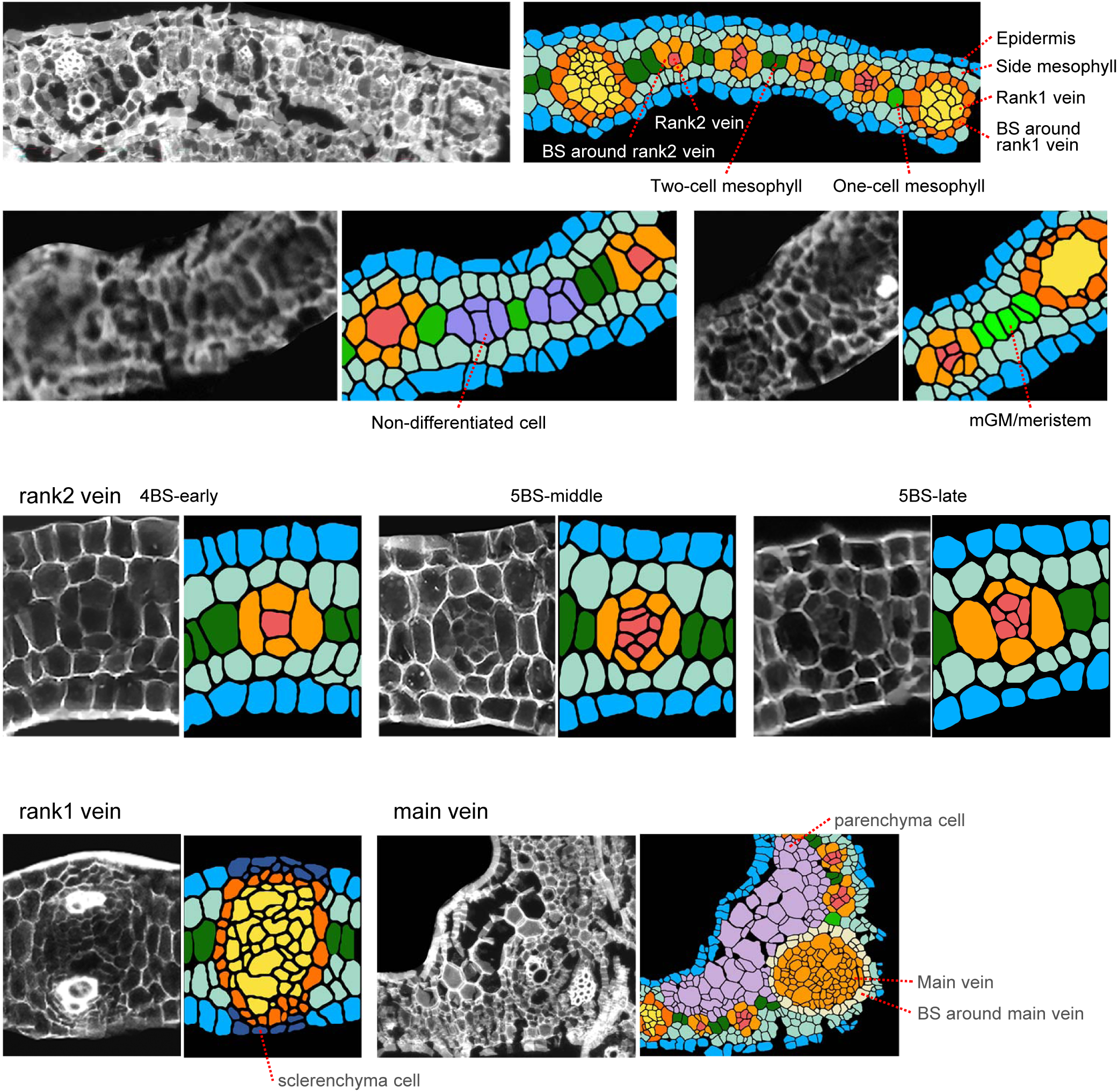
Representative picture of cell type recognized in spatial transcriptome. Left: Fluorescence-stained images; Right: Recognized cell types with artificial color labels.

**Supplementary figure 3.**
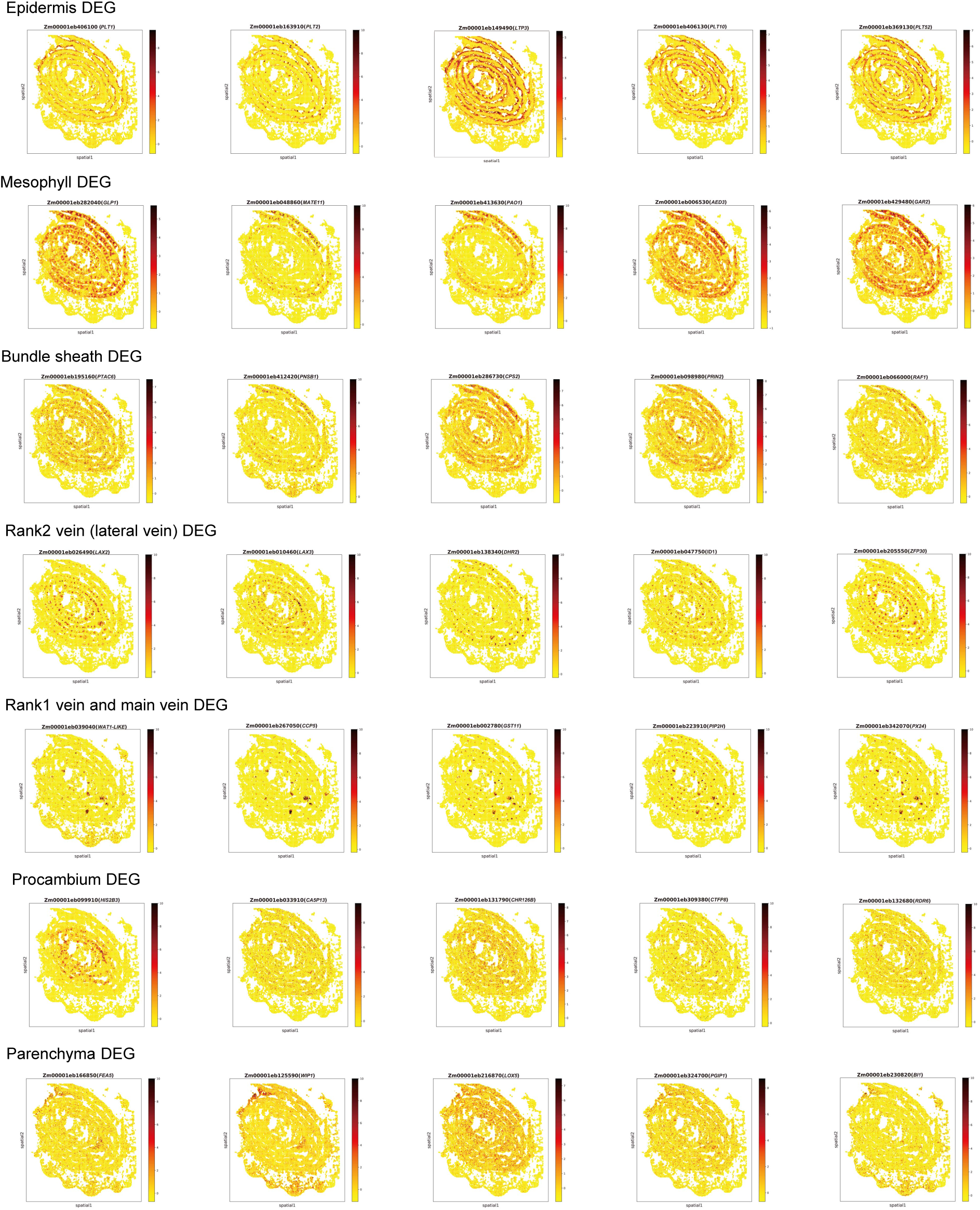
Expression patterns of marker genes identified in each cell type on Stereo-seq maps.

**Supplementary figure 4.**
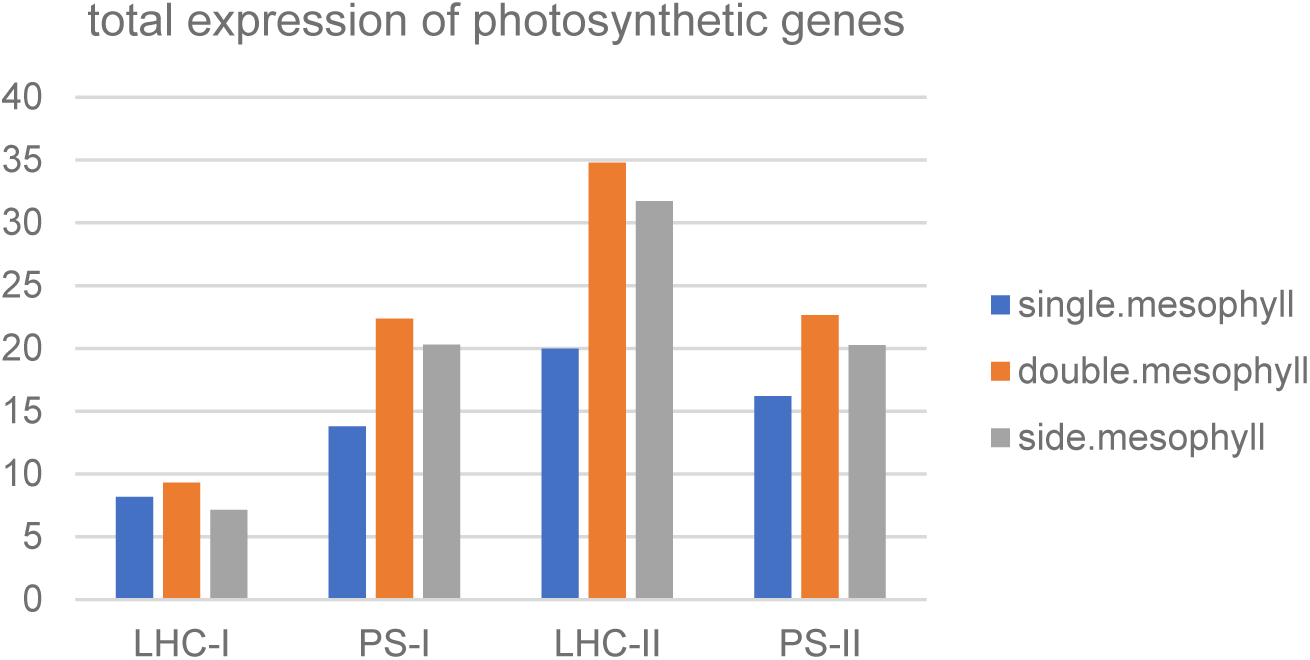
Expression patterns of photosystem related genes in three sub-types of mesophyll cell. Total expression of photosynthetic genes in in three sub-types of mesophyll cell.

**Supplementary figure 5.**
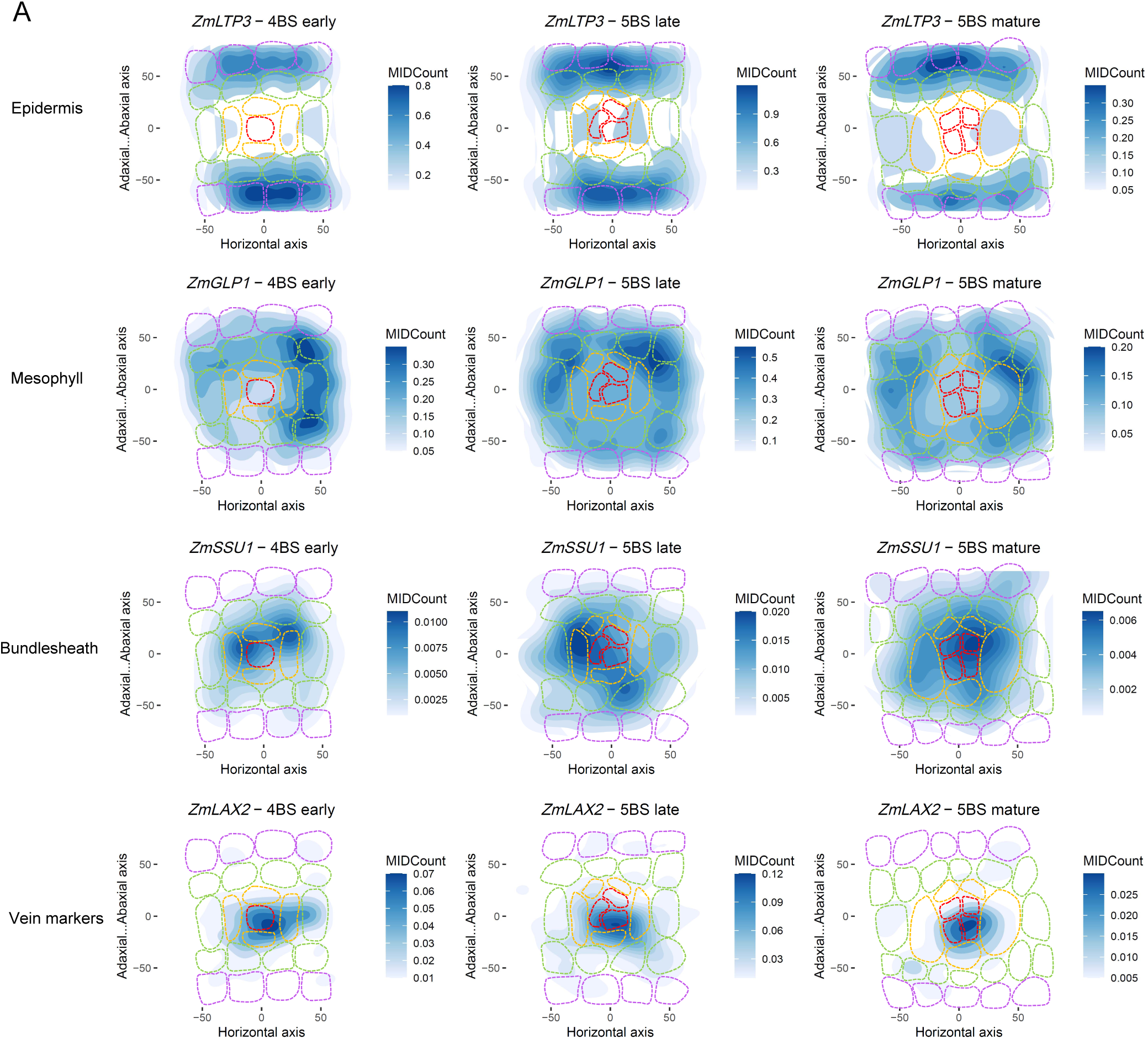
Expression patterns of marker genes of four cell types on three stages of pseudo-Kranz.

**Supplementary figure 6.**
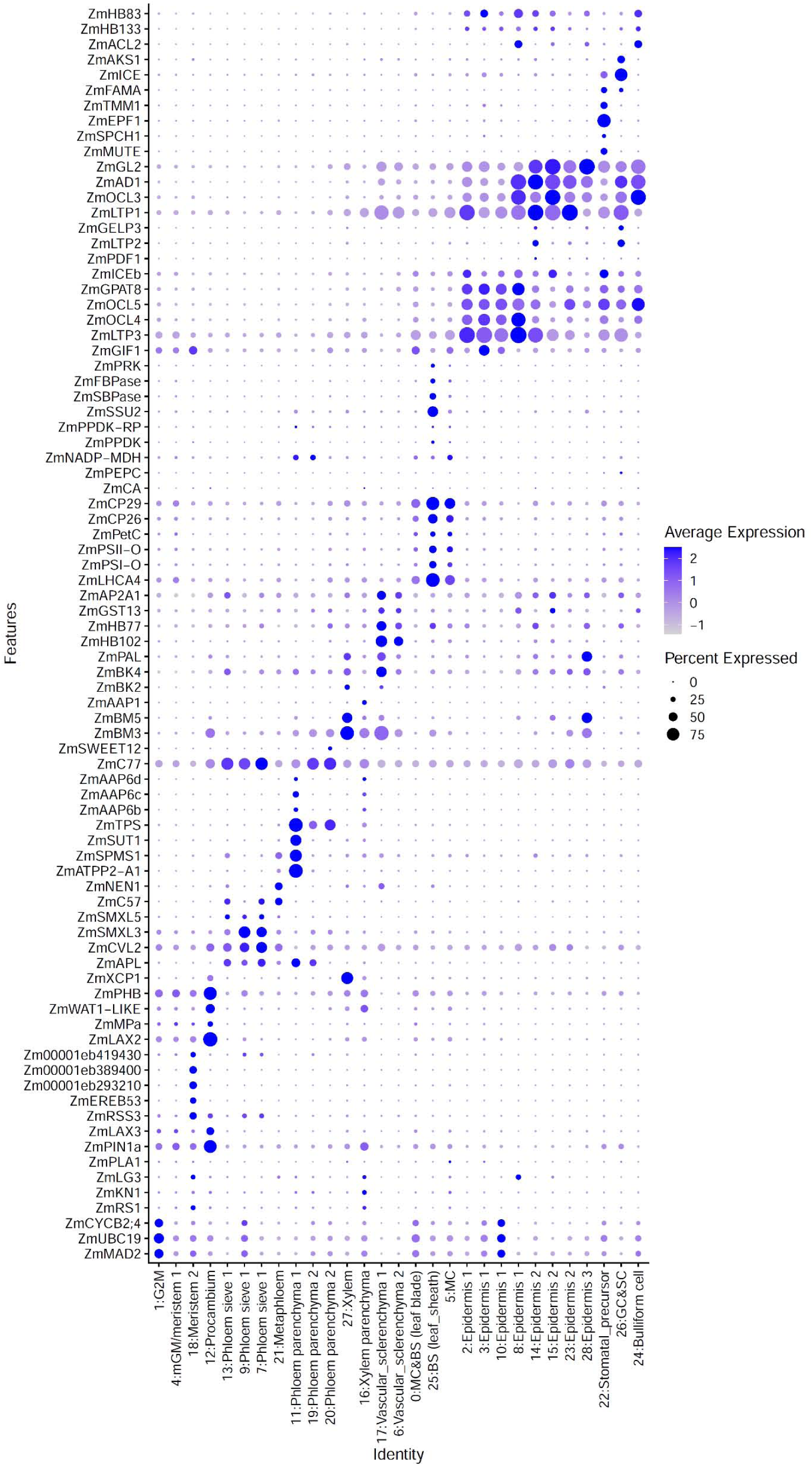
Dot plot of marker genes in the indicated cell types of maize leaf atlas. The color key from light blue to dark blue indicates low to high gene expression levels of averaged scaled data. The dot size indicates the ratio of cells with expression compared to cells in that cluster.

**Supplementary figure 7.**
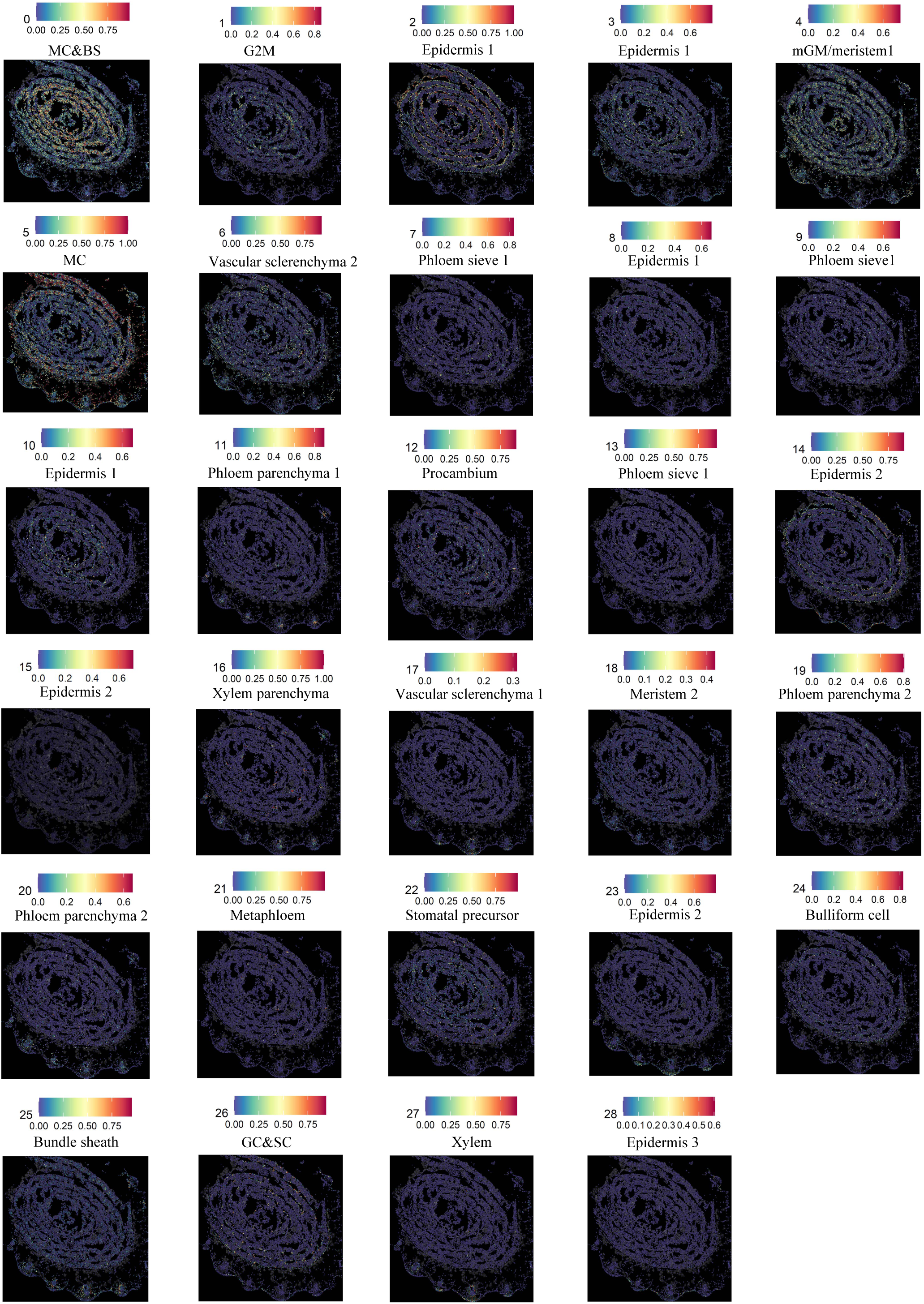
The integration of clusters of snRNA-seq atlas with cells of spatial transcriptomes of maize. The color key from blue to red indicates low to high correlation of transcriptome between the cluster and each cell of spatial atlas.

**Supplementary figure 8.**
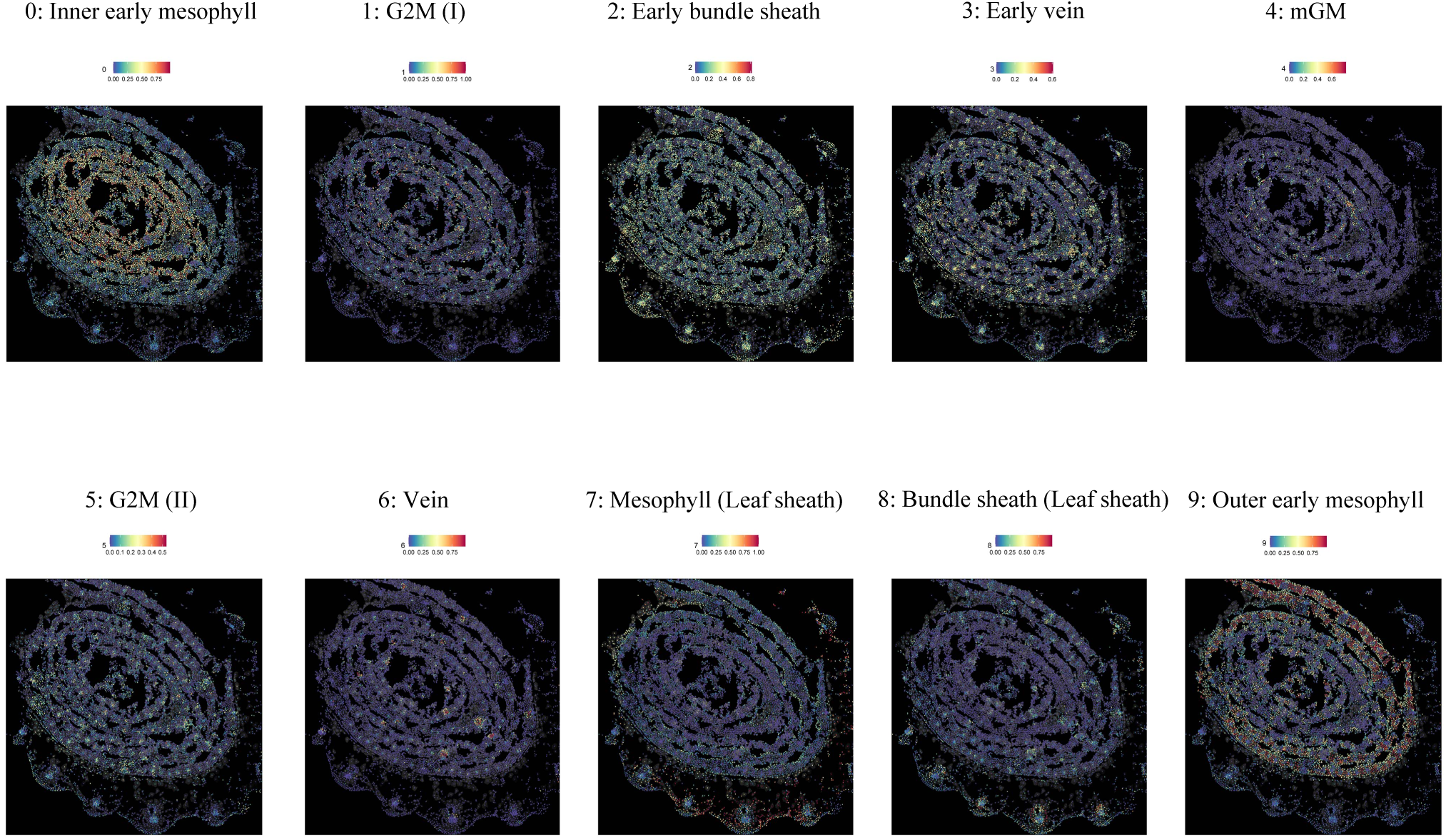
The integration of sub-clusters of snRNA data with cells of spatial transcriptomes. The color key from blue to red indicates low to high correlation of transcriptome between the cluster and each cell of spatial atlas.

**Supplementary figure 9.**
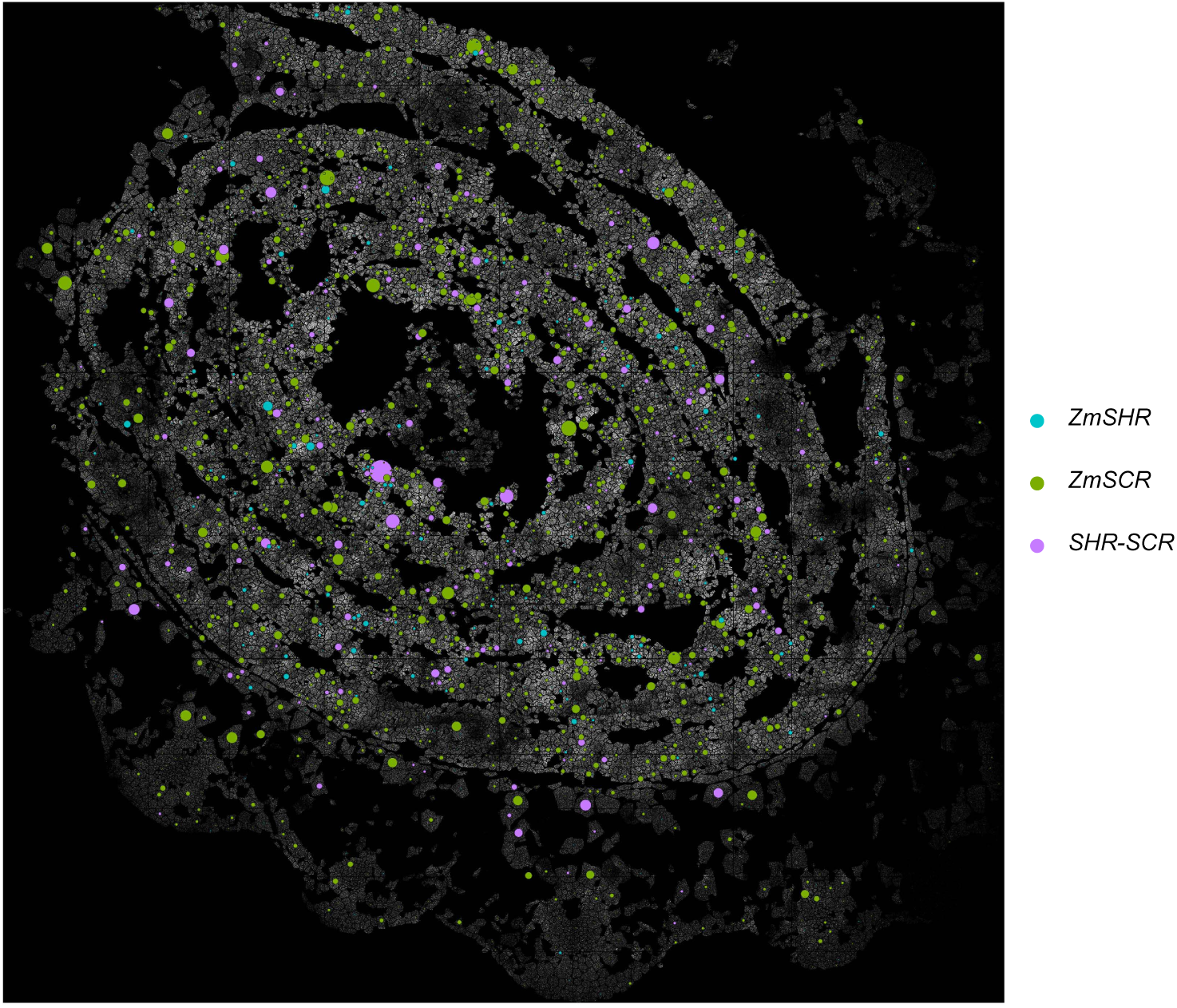
co-expression patterns of *SHR* and *SCR* genes in spatial transcriptomes. The size of the dots on the plot indicates the average expression levels of different genes. Light blue indicates the average expression of *ZmSHR1* and *ZmSHR2* genes only, green indicates the average expression of *ZmSCR1* and *ZmSCR2* genes only, and pink indicates the average expression of co-expressed *ZmSHR* and *ZmSCR* genes.

**Supplementary figure 10.**
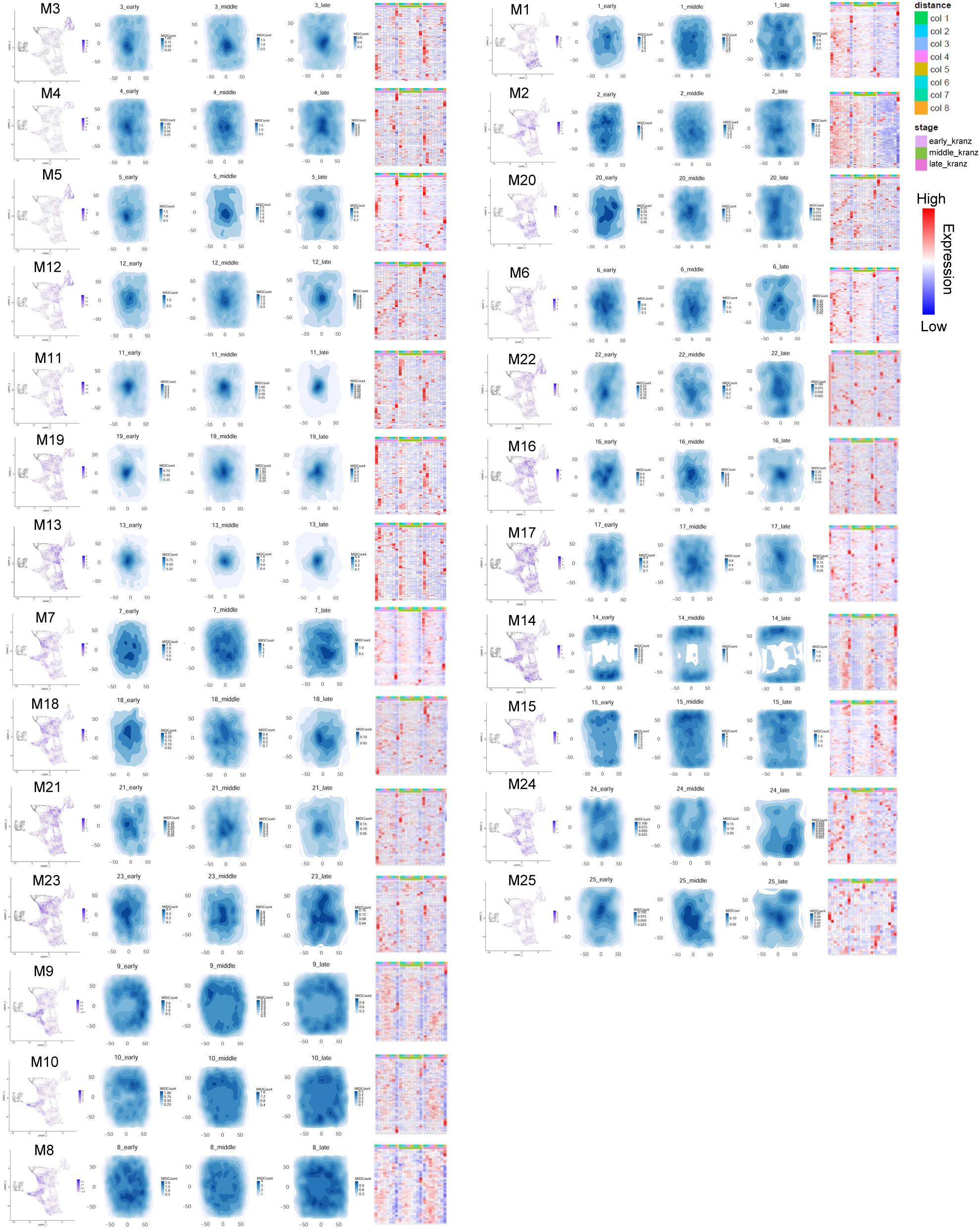
Expression patterns of module genes in snRNA-seq and pseudo-Kranz. Left : Modules scores of each cells displayed on the UMAP of subset clusters. Middle: Total expressions of gene in modules showed on pseudo-Kranz. Right: The heatmap showing the relative scaled expression of genes within each module at different distances from the center of pseudo-Kranz at three stages.

**Supplementary figure 11.**
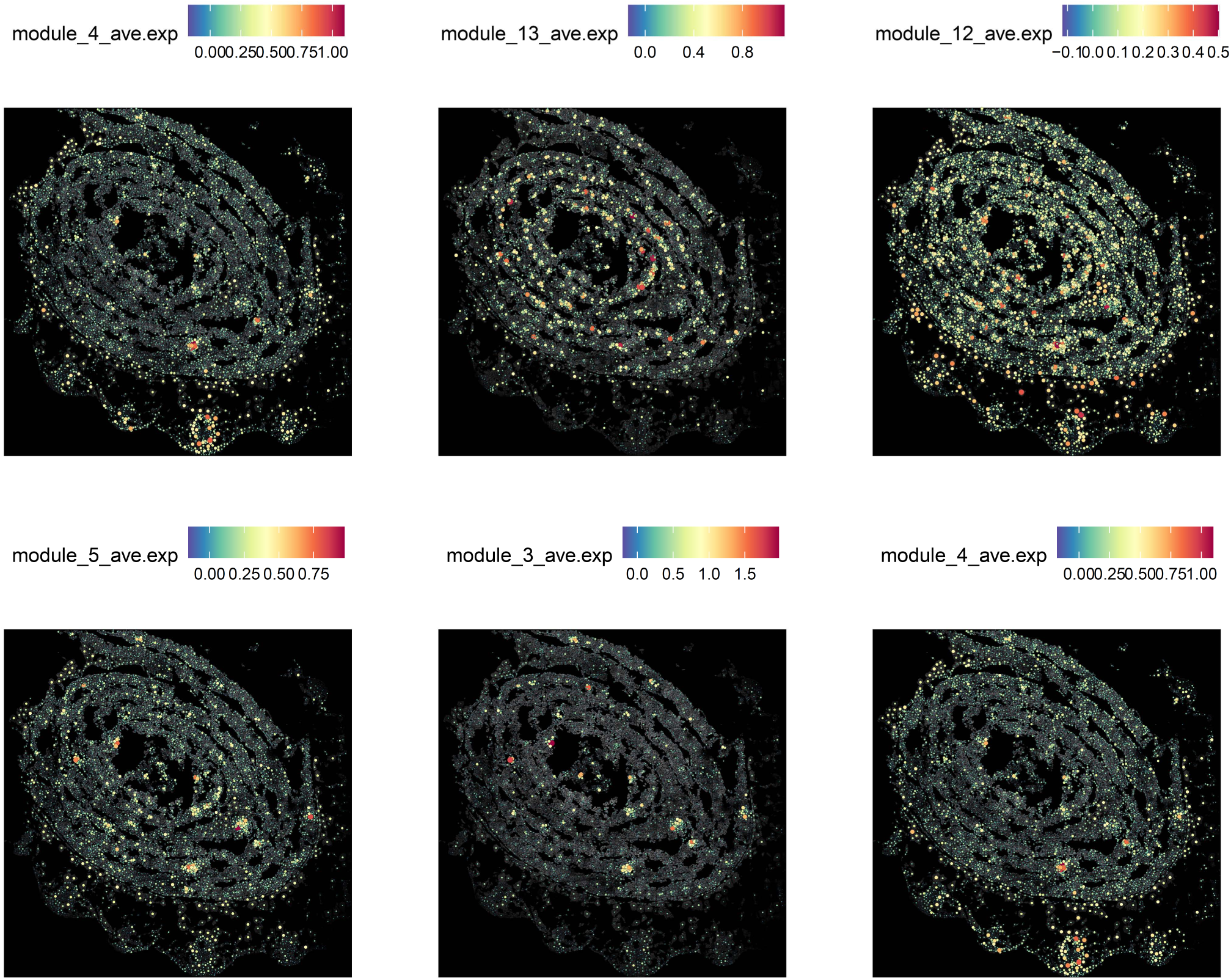
Expression patterns of individual vein-associated module genes spatial atlas.

**Supplementary figure 12.**
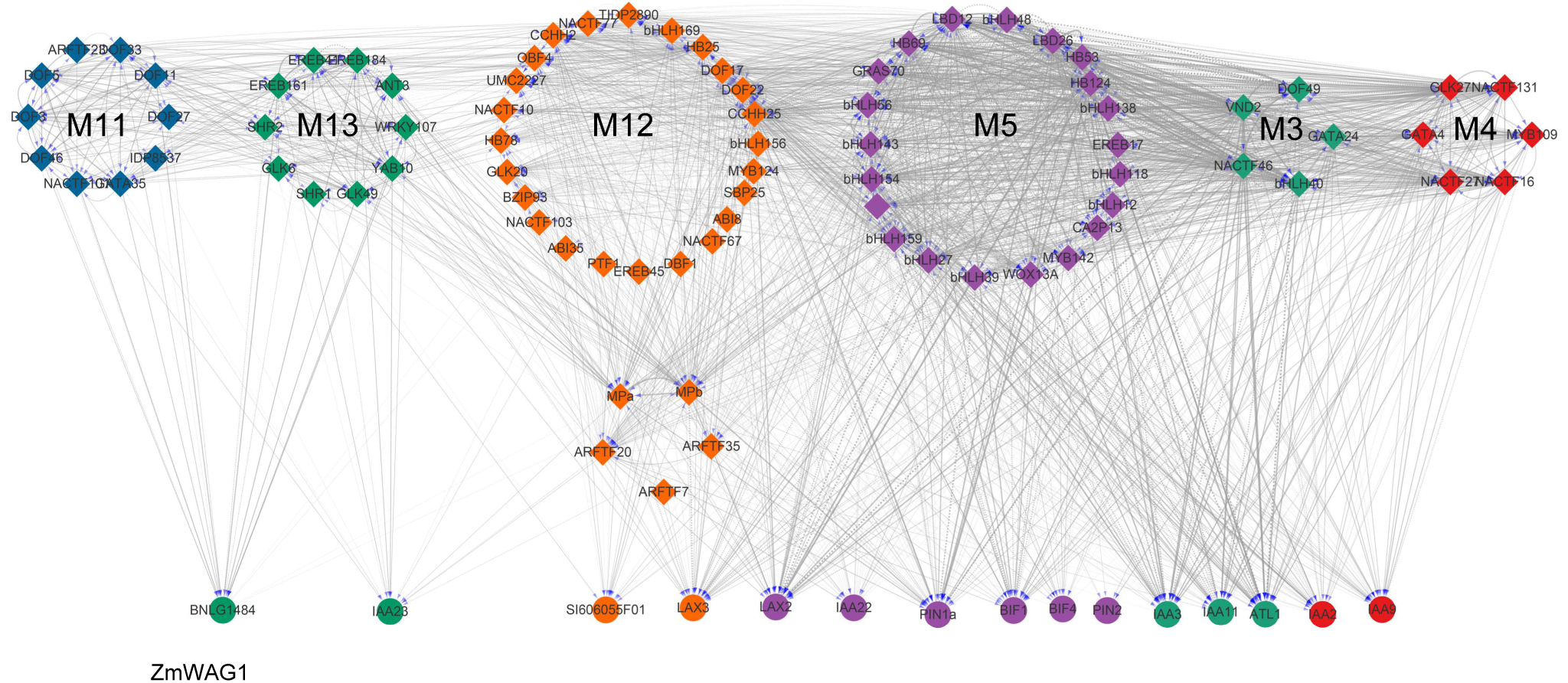
Regulatory co-expression network of vein related module genes. Colors of nodes denote distinct modules. The size of nodes reflects the aggregate correlation score. Solid lines indicate predicted regulatory interactions between TFs, while dashed lines represent potential but unconfirmed relationships without predicted binding sites. Circles and diamonds of spots represent functional genes and TFs, respectively.

**Supplementary figure 13.**
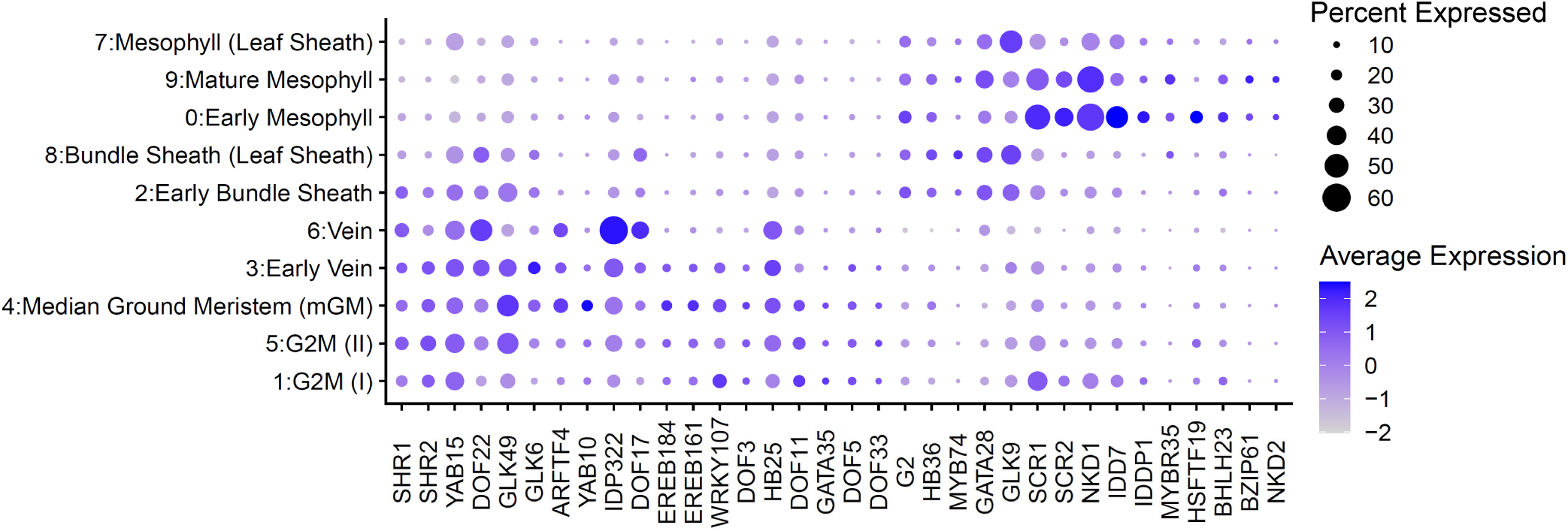
The expression patterns of candidate regulators of *SHR*, *SCR* and *G2* genes. The color key from light blue to dark blue indicates low to high gene expression levels of averaged scaled data. The dot size indicates the ratio of cells with expression compared to cells in that cluster.

**Supplementary figure 14.**
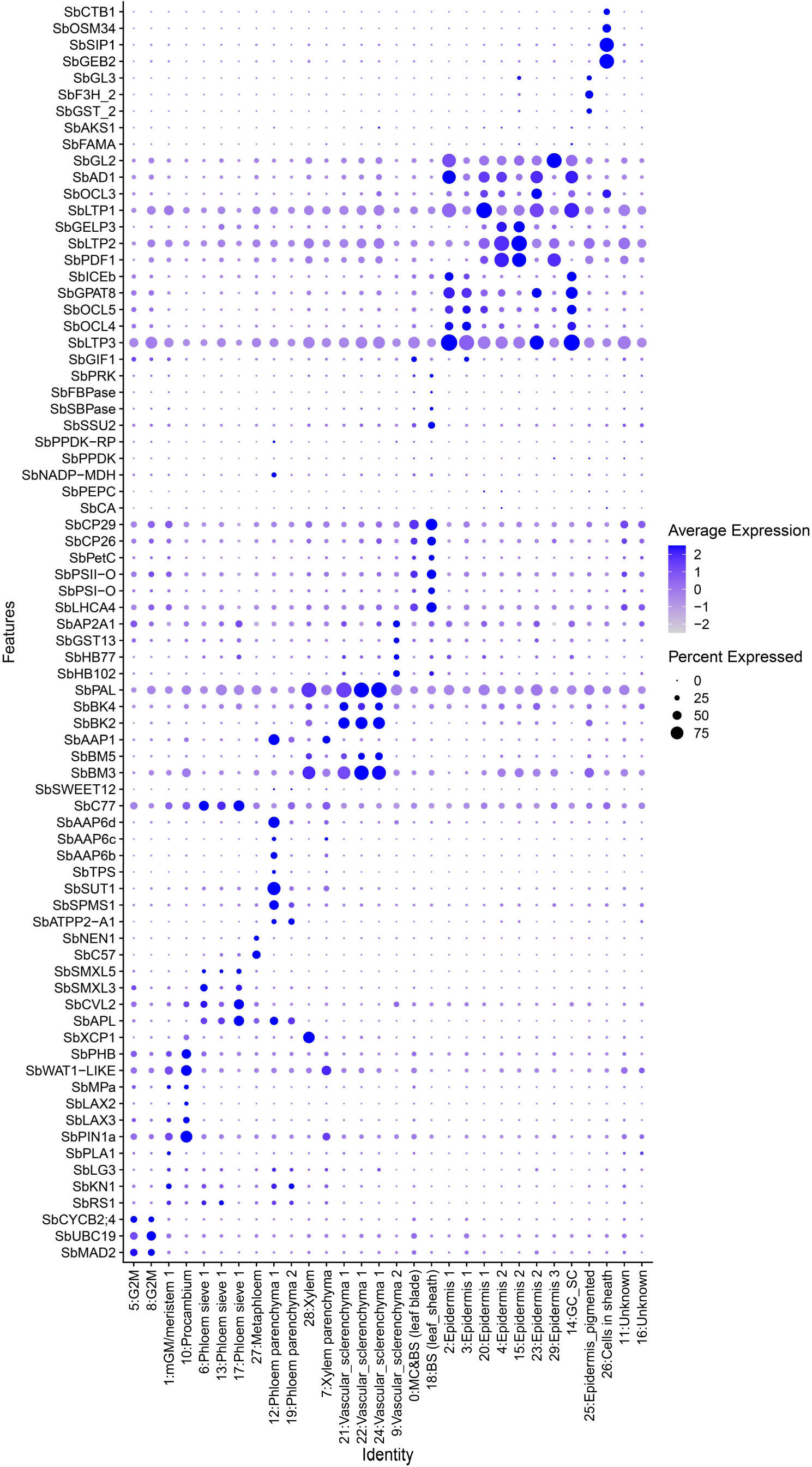
Dot plot of marker genes in the indicated cell types of Sorghum leaf atlas. The color key from light blue to dark blue indicates low to high gene expression levels of averaged scaled data. The dot size indicates the ratio of cells with expression compared to cells in that cluster.

**Supplementary figure 15.**
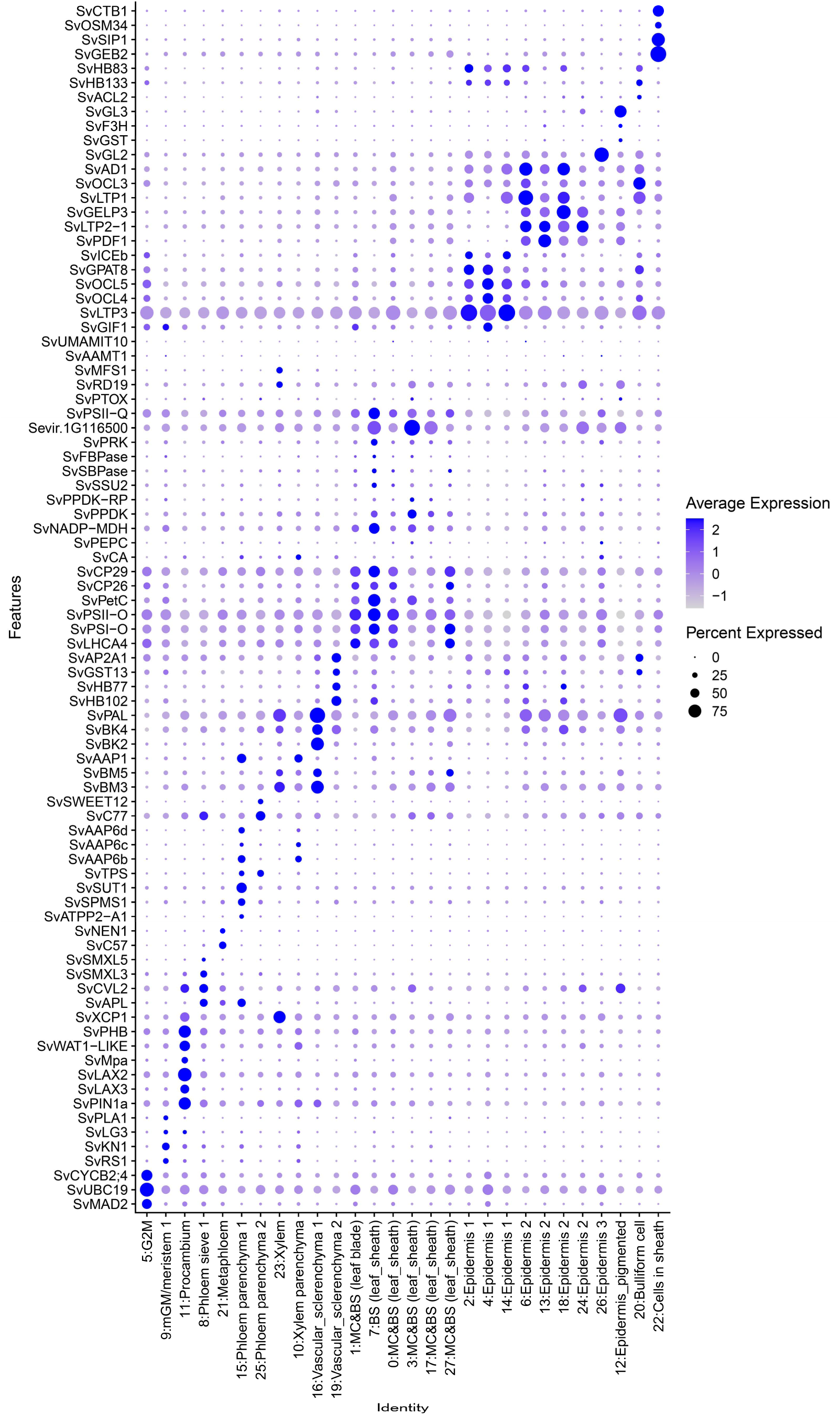
Dot plot of marker genes in the indicated cell types of Setaria leaf atlas. The color key from light blue to dark blue indicates low to high gene expression levels of averaged scaled data. The dot size indicates the ratio of cells with expression compared to cells in that cluster.

**Supplementary figure 16.**
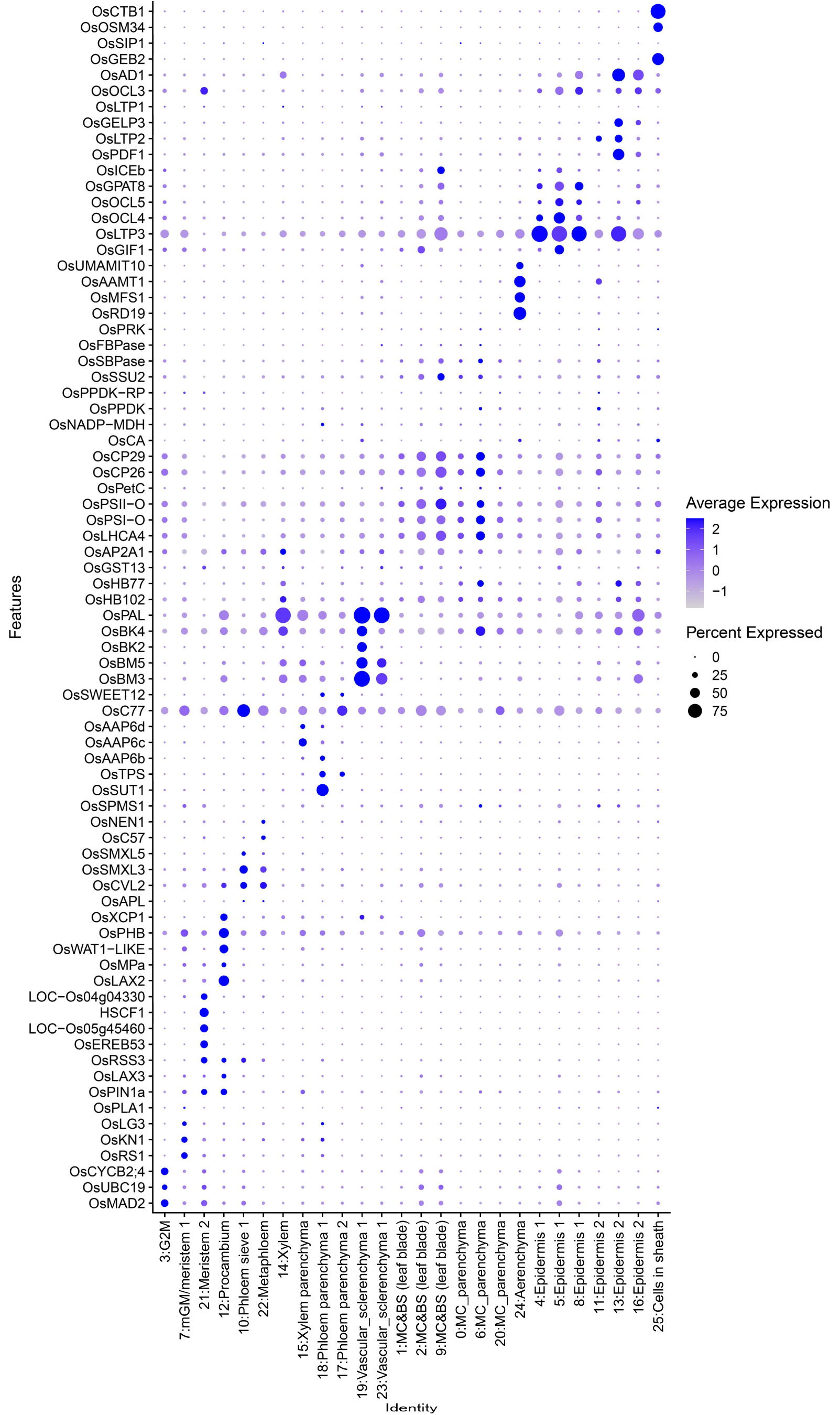
Dot plot of marker genes in the indicated cell types of rice leaf atlas. The color key from light blue to dark blue indicates low to high gene expression levels of averaged scaled data. The dot size indicates the ratio of cells with expression compared to cells in that cluster.

**Supplementary figure 17.**
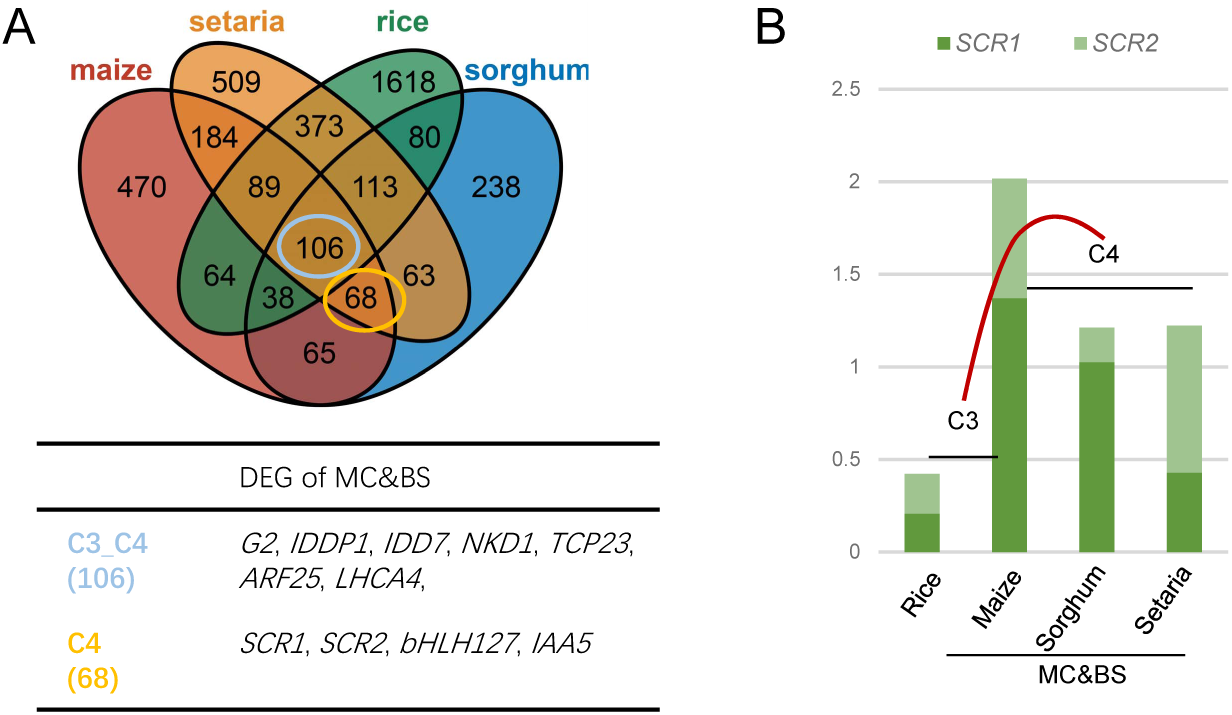
Conserved DEGs in BSC and/or MC among C_4_ and C_3_ plants. (A) Comparison of DEGs in clusters of MC&BSC between C_4_ and C_3_ species. Up: Venn diagrams showing the numbers of DEGs in different species; Bottom: C_4_ specific or C_4_ and C_3_ shared representative markers are listed in table. The relative expression levels of *SCR* genes in clusters of MC&BSC of four species

**Supplementary figure 18.**
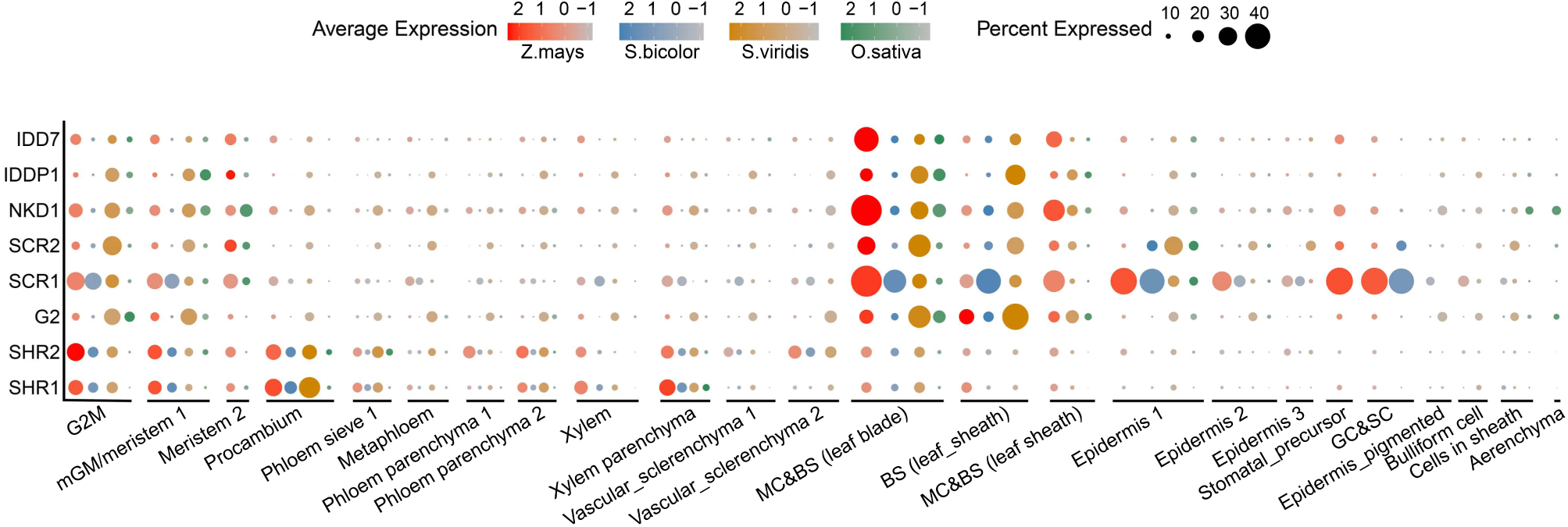
The expression patterns of selected key genes in different cell types of C_4_ and C_3_ plant. The color key (red for maize, blue for Sorghum, brown for Setaria and green for rice) from light to dark indicates low to high gene expression levels of averaged scaled data. The dot size indicates the ratio of cells with expression compared to cells in that cluster.

**Supplementary figure 19.**
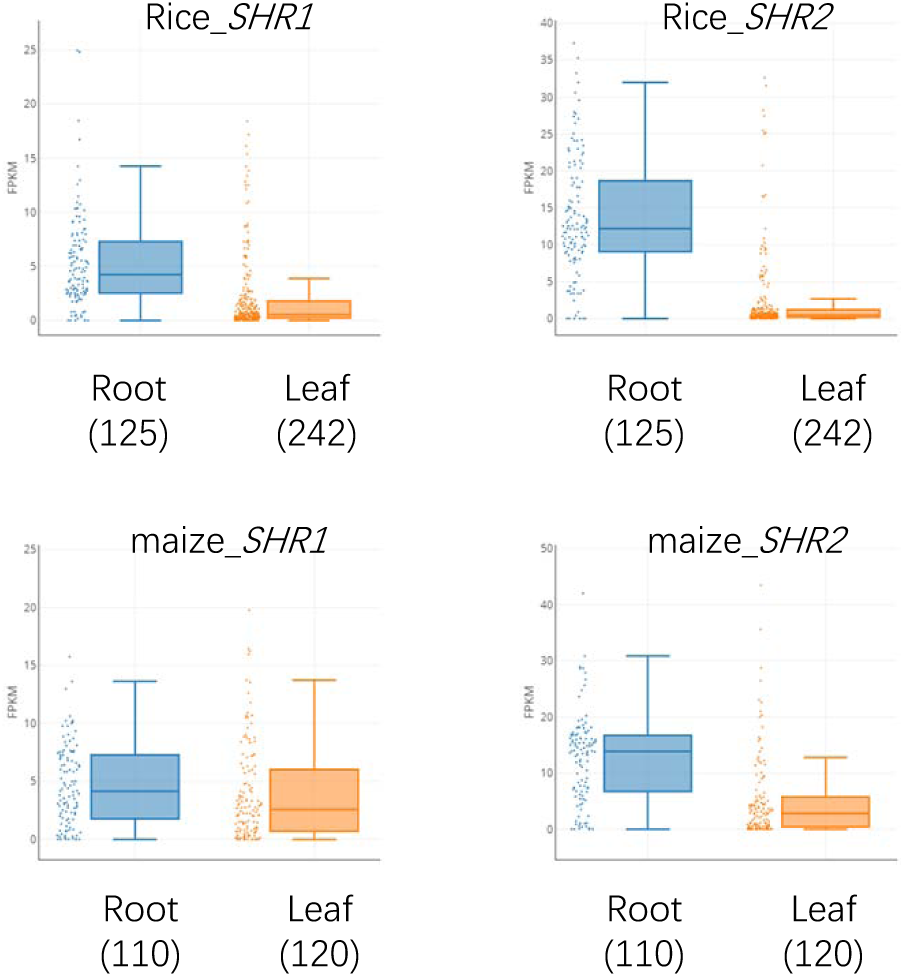
The expression levels of *SHR* genes in root and leaf of maize and rice.

**Supplementary figure 20.**
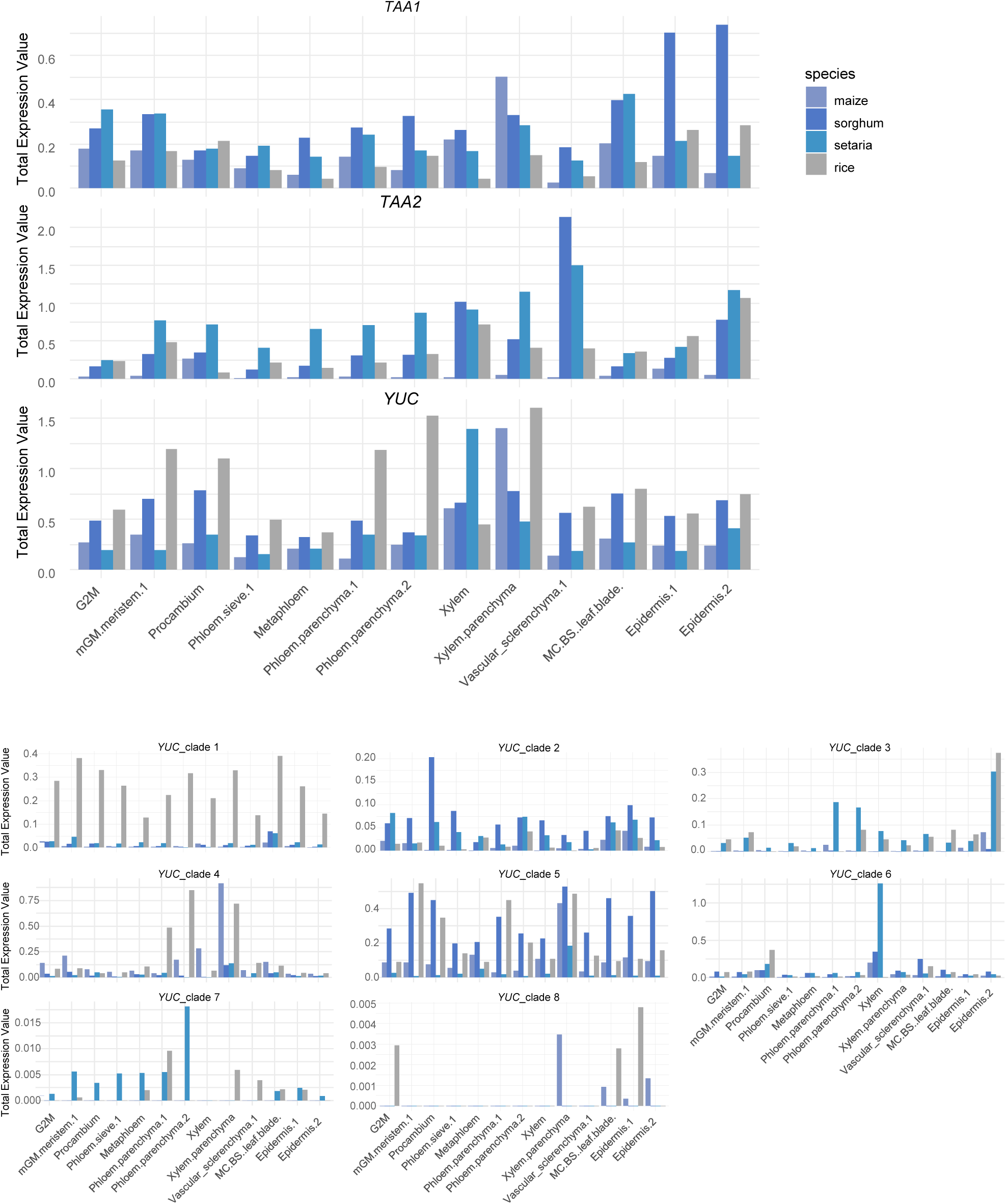
The expression levels of auxin biosynthesis genes in C_4_ and C_3_ species.

**Supplementary figure 21.**
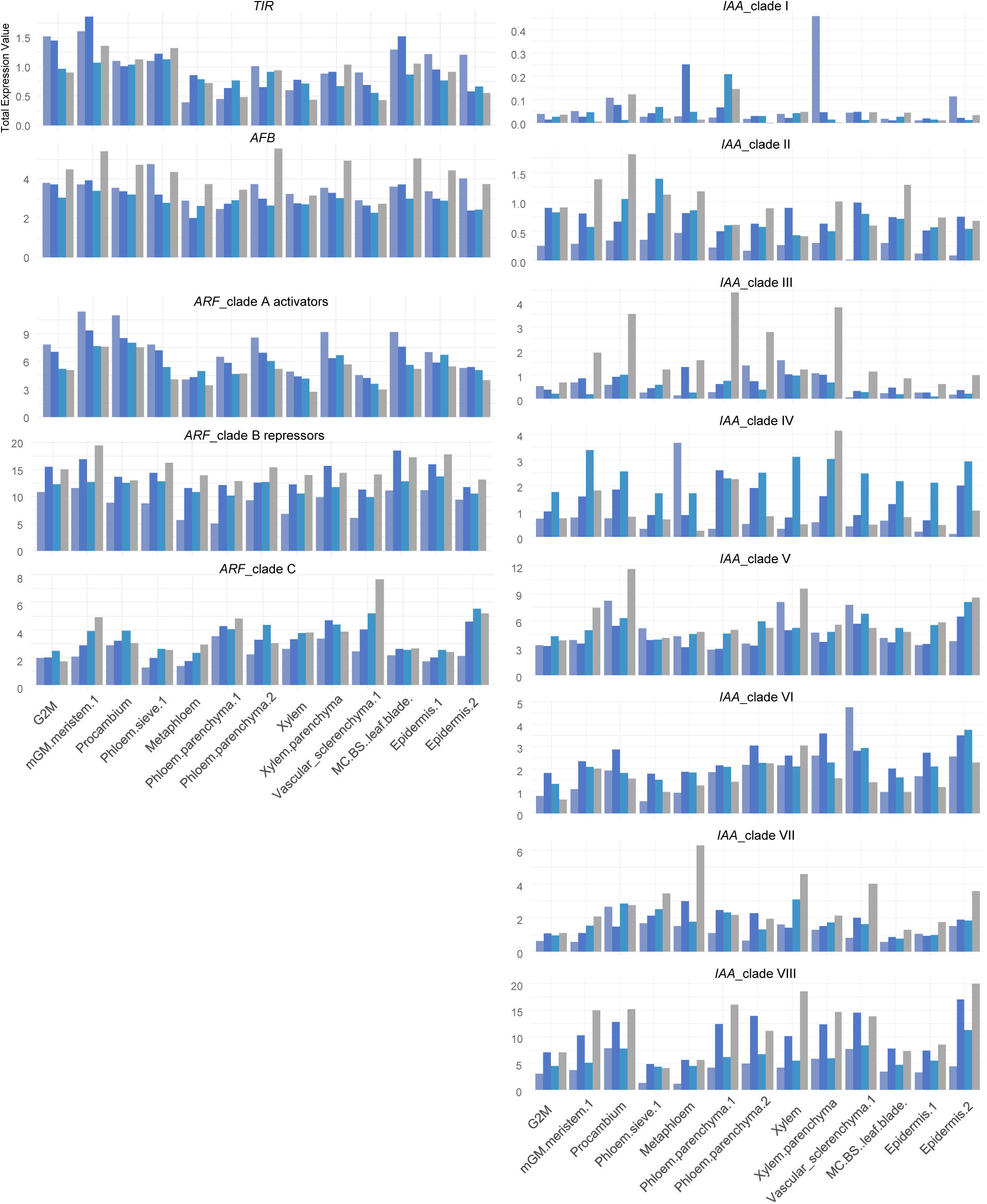
The expression levels of auxin signal transduction genes in C_4_ and C_3_ species.

**Supplementary figure 22.**
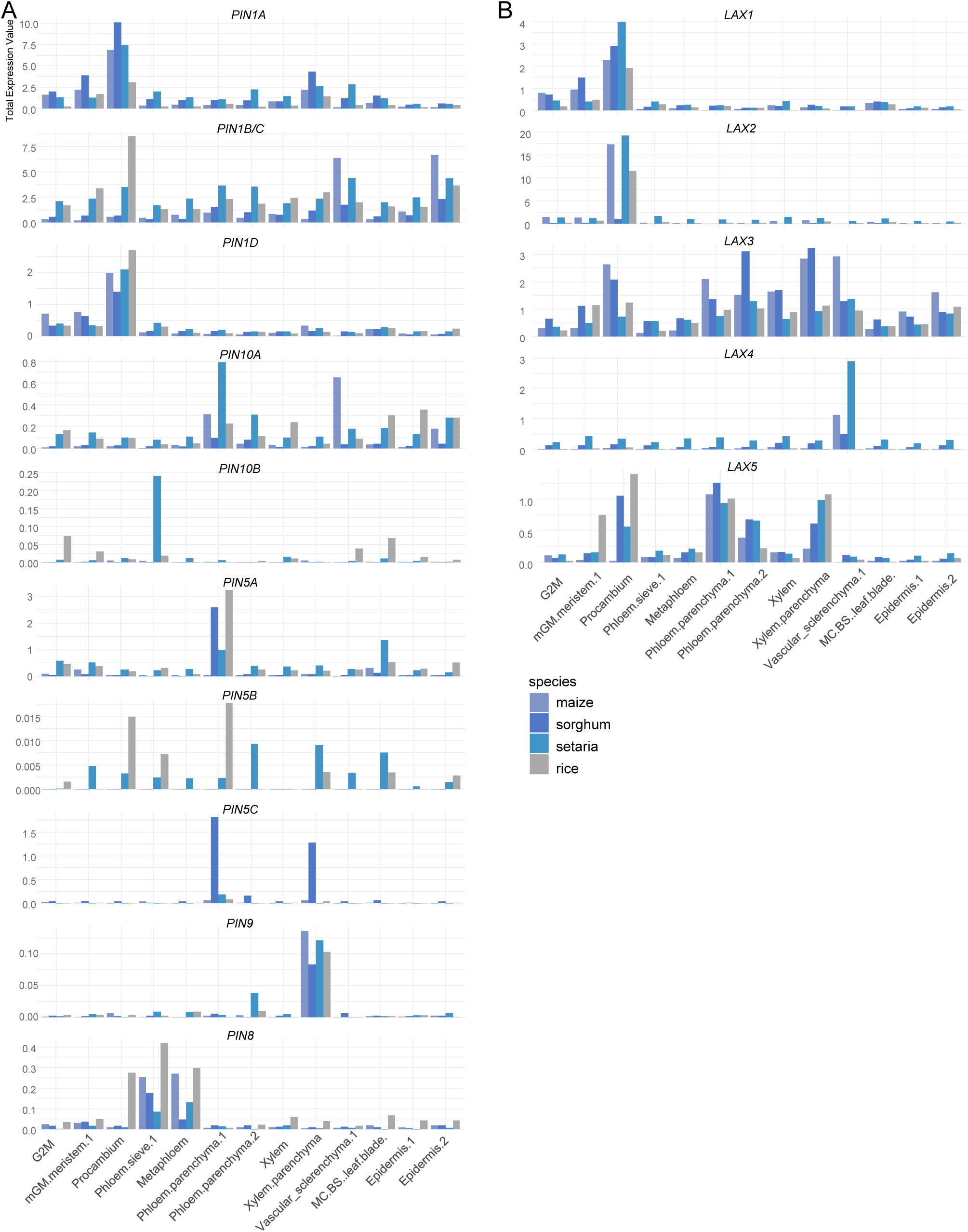
The expression levels of auxin transport genes in C_4_ and C_3_ species.

**Supplementary figure 23.**
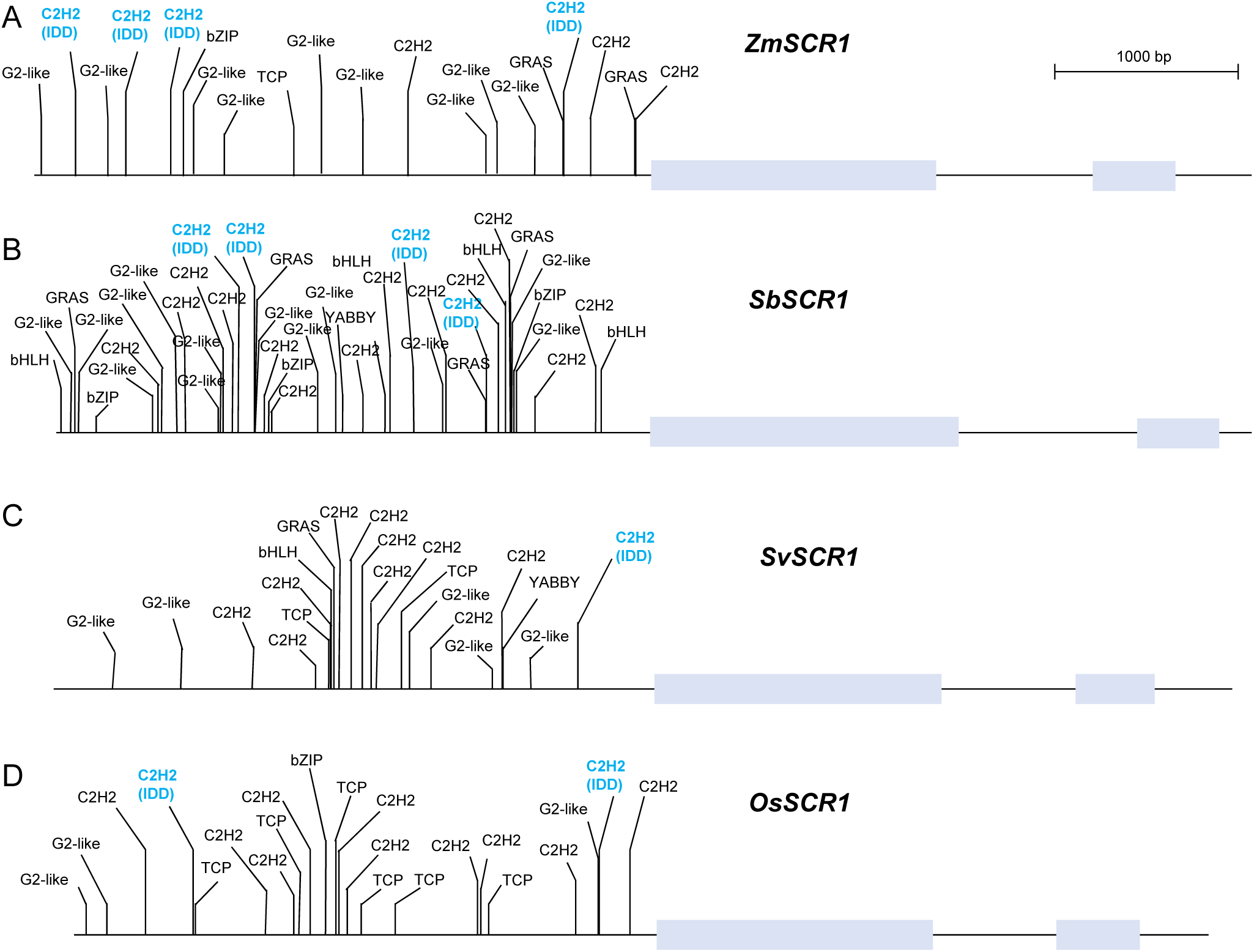
The relative positions of predicted binding sites of TFs co-expressed with *SCR1*. The transcription factors binding sites (TFBS) located approximately 3 kb upstream of the coding sequence in maize (A), sorghum (B), Setaria (C), and rice (D). The horizontal lines represent gene lengths, and the boxes on these lines indicate the positions of the two exons within the SCR1 genes. The vertical lines mark the relative positions of TFBS, with the corresponding transcription factor families labeled above each vertical line. IDD binding sites are highlighted in blue.

**Supplementary figure 24.**
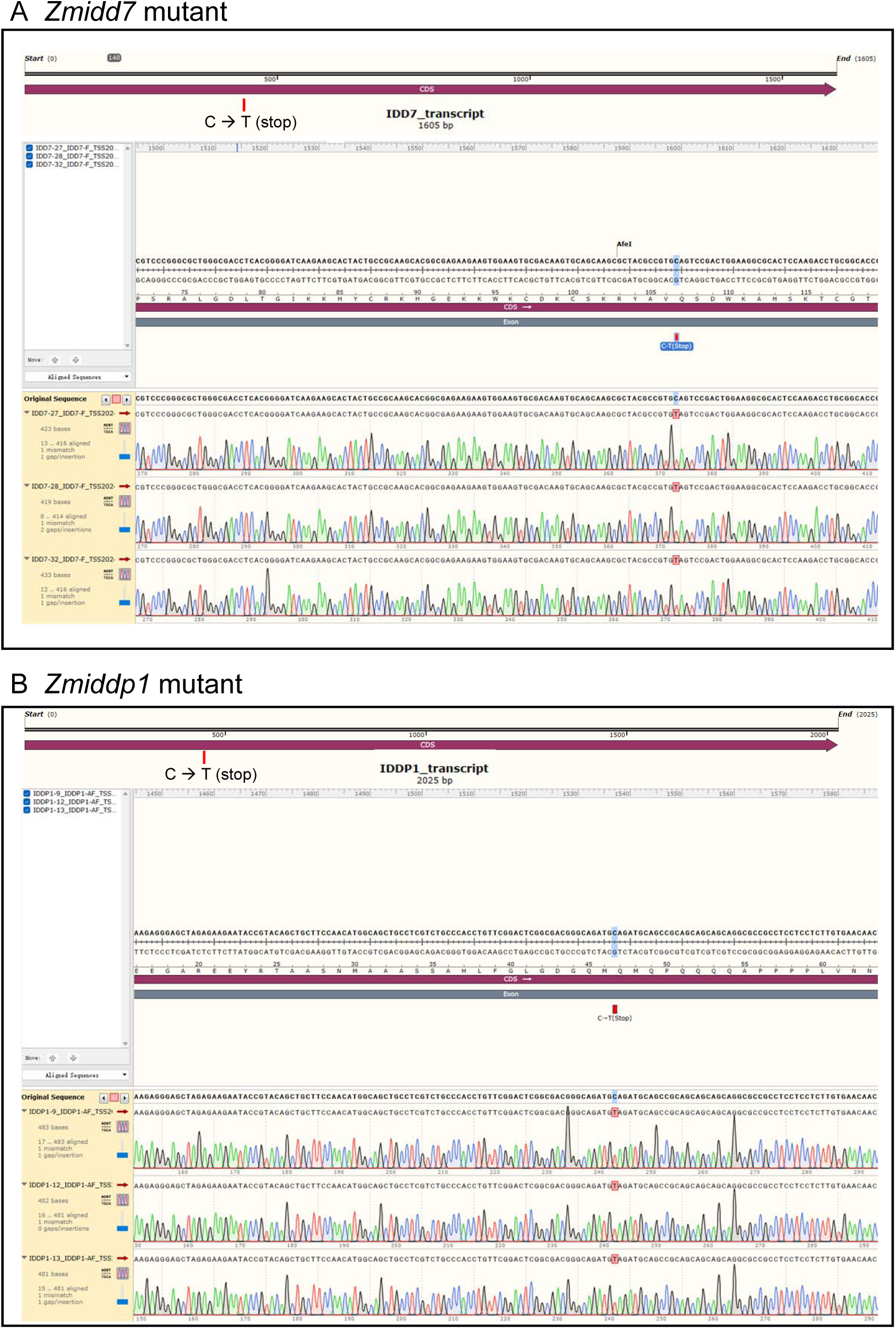
The mutation site identification of EMS mutants. (A) There is a mutation of C to T at the 481^st^ bp among full-length 1605bp coding sequence of *ZmIDD7*, which makes the 161^st^ amino-acid glutamine (Q, CAG) change to a stop codon (TAG) that leads a premature termination on translation of full-length 534 amino-acids. (B) There is a mutation of C to T at the 448^th^ bp among full-length 2025bp coding sequence of *ZmIDDP1*, which makes the 150^th^ amino-acid glutamine (Q, CAG) change to a stop codon (TAG) that leads a premature termination on translation of full-length 674 amino-acids.

